# Toll-like receptor 2 orchestrates a potent anti-tumor response in non-small cell lung cancer

**DOI:** 10.1101/2021.06.04.446876

**Authors:** Fraser R. Millar, Adam Pennycuick, Morwenna Muir, Andrea Quintanella, Priya Hari, Elisabeth Freyer, Philippe Gautier, Alison Meynert, Vitor H. Teixeira, John Connelly, William AH Wallace, Andrew H Sims, Margaret C. Frame, Luke Boulter, Sam M. Janes, Simon Wilkinson, Juan Carlos Acosta

**Author notes:** **Corresponding author: Dr Juan-Carlos Acosta, Address:** Cancer Research UK Edinburgh Centre, Institute of Genetics and Cancer, University of Edinburgh, Crewe Road South, Edinburgh, EH4 2XU, UK, **Email:**.

## Abstract

Targeting early-stage lung cancer is vital to improve overall survival. We previously identified Toll-like receptor 2 (TLR2) as a regulator of oncogene-induced senescence (OIS) and the senescence-associated secretory phenotype (SASP), both key for tumor suppression. Here, we demonstrate that TLR2 is widely expressed in human lung tumor epithelium where it correlates with improved survival and clinical regression. Using genetically engineered mouse models of lung cancer we have shown that *Tlr2* is a tumor suppressor in lung cancer initiation via regulation of proliferation and the SASP. The SASP is integral in the regulation of immune surveillance of premalignant cells, and we observe impaired myeloid derived immune surveillance following *Tlr2* loss. Lastly, we show that administration of a synthetic Tlr2 agonist significantly reduces preinvasive lung tumor growth. Our data highlight an unexpected tumor surveillance pathway in early-stage lung cancer with therapeutic potential.

**Statement of significance:** Lung cancer is a major cancer of unmet need. This study identifies a novel tumor suppressor mechanism in lung cancer. Not only does this highlight a potential therapeutic target for early-stage disease but also multiple secreted candidate biomarkers that could be exploited to augment lung cancer screening approaches.

## Introduction

Lung cancer is the most lethal cancer type worldwide, with a mortality rate greater than breast, colorectal and prostate cancer combined (1). Non-small cell lung cancer (NSCLC) accounts for 88% of cases (2), with lung adenocarcinoma (LUAD) and lung squamous carcinoma (LUSC) the predominant histological subtypes. Early-stage NSCLC is associated with a favourable 5-year survival rate of over 50% (3). However, the majority of patients present with advanced-stage disease which has an abysmal 5-year survival rate of just 6% (3). Recent advances with immunotherapy and targeted therapies against identifiable driver oncogenes has improved outcomes in a small subset of NSCLC patients (4–6). However, due to the low frequency of targetable mutations and the inevitable emergence of resistance and disease relapse, overall outcomes are still very poor. In order to improve lung cancer survival, it is clear that we must understand and target early-stage disease. Indeed, two large multicentre randomised clinical trials have demonstrated a clear mortality benefit with the use of low dose computed tomography (LDCT) screening in patients at high risk of lung cancer (7,8). This approach is not without its drawbacks; LDCT screening has a high false positive rate, resulting in significant anxiety and potentially dangerous investigations in healthy patients. Furthermore, risk prediction tools used to assess an individual’s suitability for screening can underestimate risk in certain patient demographics (9) and it has been reported that only one third of new lung cancer patients would have met eligibility criteria for LDCT screening (10). Thus, a further understanding of the biology of early-stage disease may open up new therapeutic avenues and highlight novel ways to identify disease earlier. Oncogene-induced senescence (OIS) is a terminal stress response that is initiated following the activation of oncogenes and is a key tumor suppressor mechanism (11). Along with a robust cell cycle arrest mediated by p53-p21 and p16-Rb tumor suppressor pathways (12), OIS is characterised by significant metabolic upregulation and expression of a variety of secreted factors collectively termed the senescence-associated secretory phenotype (SASP) (13,14). Regulation of the SASP is complex and context dependent. While many stressors that induce the SASP also induce the growth arrest (for example DNA damage), the SASP can be induced independently of tumor suppressor pathways (14–16). Nonetheless, the SASP contributes to the non-cell autonomous tumor suppressor effects of OIS via reinforcing the senescence-associated growth arrest in an autocrine manner (13), inducing senescence in neighbouring cells (paracrine senescence) (17) and orchestrating immune cell mediated clearance of senescence cells (senescence surveillance) (18,19). On the other hand, the SASP can also induce a permissive environment accelerating cancer progression (20). Thus, elucidating the mechanisms controlling the tumor suppressive and promoting functions of the SASP, and identifying which specific factors play a functional role in cancer progression is key to successfully manipulate senescence in anti-cancer therapies. Markers of OIS are widespread in preinvasive tumors in the lung, however this expression is lost during the progression to invasive malignancy (21) suggesting a tumor suppressor role for OIS in lung cancer. We recently identified the innate immune receptor Toll-like receptor 2 (TLR2) as a key regulator of OIS and expression of the SASP, signalling via acute-phase serum amyloid A (A-SAA) proteins (22). However, the role of TLR2 in lung tumor progression has not yet been investigated. Here, using clinical human samples and genetically engineered mouse models (GEMMs), we have demonstrated that *TLR2* coordinates the induction of cell autonomous and non-cell autonomous tumor suppressor responses which together impair NSCLC progression. Understanding this process not only identifies novel therapeutic targets for early-stage disease but also highlights multiple candidate biomarkers to aid in screening population selection.

## Results

### TLR2 expression correlates with improved survival and clinical regression in human lung cancer

We first analysed *TLR2* expression in human LUAD samples from the cancer genome atlas (TCGA) (23) and found that *TLR2* expression significantly correlated with improved survival (**Figure 1A**). This correlation was specific to LUAD as *TLR2* expression had no such correlation in other cancer types (**Supp figure 1A-C**). Moreover, this correlation in LUAD was also observed in genes encoding TLRs that form heterodimers with TLR2 upon activation (TLR1, 6, 10) but not with the other plasma membrane toll-like receptors (such as TLR4) (**Supp figure 1D-F**), indicating a specific role for the TLR2 sensing networks in LUAD. To test which cellular compartment *TLR2* is expressed in, we analysed histological samples of human LUAD and performed immunohistochemistry (IHC) staining for TLR2. Strikingly, we found that TLR2 expression is significantly increased in LUAD tumor epithelium in comparison to paired normal epithelium (**Figure 1C-D**). To further investigate the role of *TLR2* signalling in early and late stage LUAD we analysed a published RNA sequencing dataset from a cohort of preinvasive (adenocarcinoma in situ – AIS, and minimally invasive adenocarcinoma – MIA) and invasive LUAD samples (24). The gene expression profiles of AIS and MIA lesions were similar (**Supp figure 1G**); hence we grouped these lesion types together for further analysis. *TLR2* gene expression was significantly increased in preinvasive tumors (AIS/MIA) compared to invasive tumors (LUAD) (**Figure 1B**), suggesting that TLR2 is induced early in lung cancer progression. However, given that these lesions are surgical resections specimens, we cannot glean any prognostic insight based on their *TLR2* expression profiles. To answers these questions requires longitudinal sampling and careful clinical follow up of individual lesions, which is not possible in surgically resected LUAD precursors. Due to their proximal location making them accessible to bronchoscopic sampling, human preinvasive LUSC lesions have undergone extensive longitudinal molecular characterisation in comparison to distally located preinvasive LUAD lesions. Longitudinal follow-up of preinvasive LUSC lesions using autofluorescence bronchoscopy has shown that not all preinvasive lesions progress to invasive malignancy. In fact, only half progress to cancer and up to 30% regress to normal epithelium (**Figure 1E**) (25). We therefore analysed transcriptomic data from a unique cohort of preinvasive LUSC samples that have been carefully clinically phenotyped, detailing either subsequent progression to cancer or regression to normal epithelium (26). We analysed the expression profiles of these samples and found that *TLR2* expression significantly correlated with subsequent clinical regression (**Figure 1F**). Importantly, these expression profiles represent lesion epithelium only as stroma was removed by laser capture microdissection prior to RNA extraction. Furthermore, the epithelial specificity of *TLR2* expression was confirmed by performing TLR2 IHC staining (**Figure 1G, Supp figure 1H**). Taken together, these data demonstrate that *TLR2* is widely expressed in human NSCLC epithelium and correlates with improved survival and clinical regression, strongly suggesting a tumor suppressor function for *TLR2* in NSCLC.

**Figure 1:**
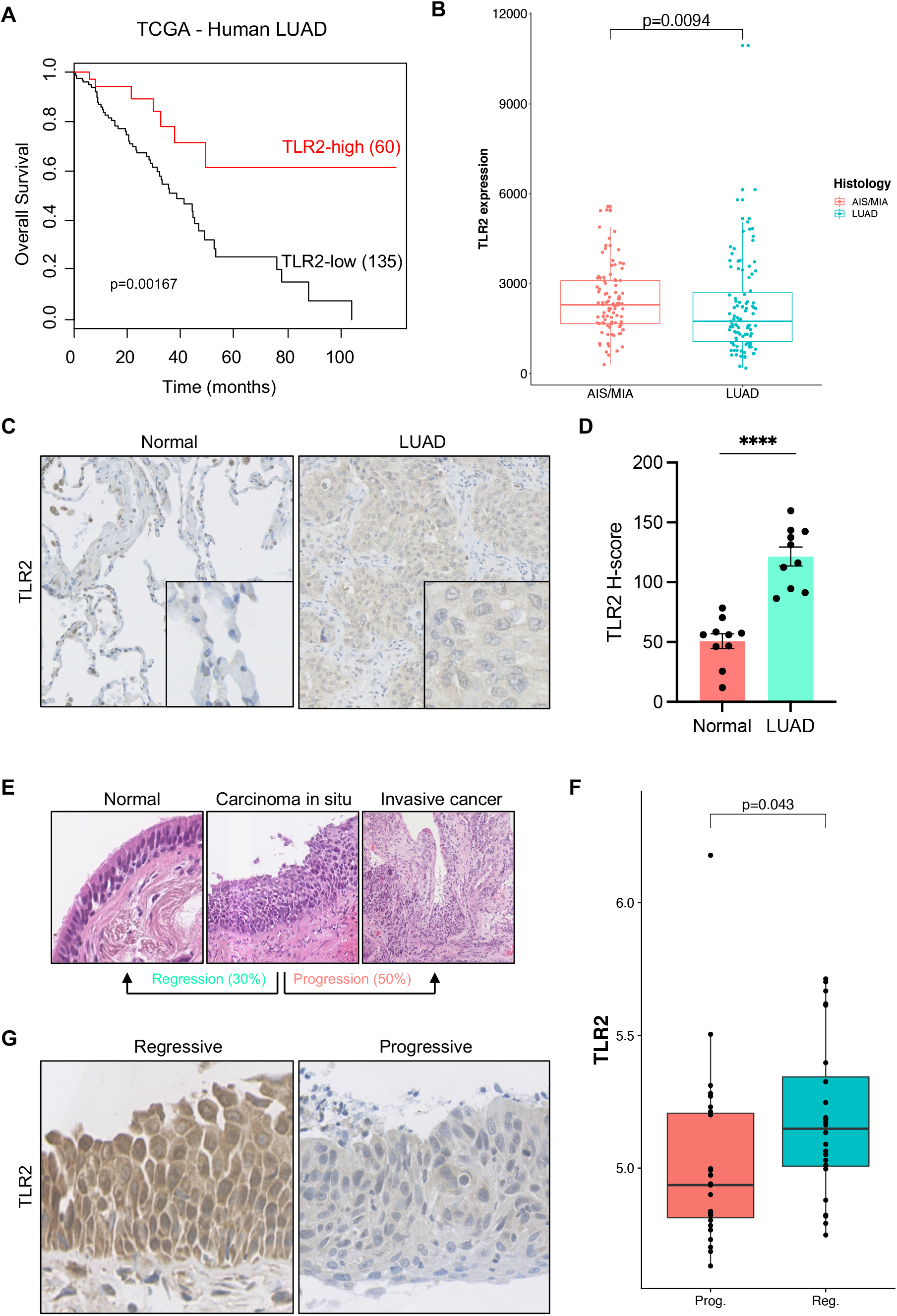
TLR2 expression correlates with improved survival and clinical regression in human lung cancer. **A,** Kaplan-Meier survival analysis of human LUAD patients based on high versus low *TLR2* expression from the cancer genome atlas (TCGA). Statistical analysis was performed using the log-rank (Mantel-Cox) test. **B,** *TLR2* gene expression was compared between LUAD precursor lesions (AIS – adenocarcinoma in situ, MIA – minimally invasive adenocarcinoma) and invasive lung adenocarcinoma lesions (LUAD). Statistical analysis was performed using the Mann-Whitney test. **C,** Representative IHC staining for TLR2 in paired normal tissue and human LUAD samples, with corresponding H-score quantification in **D**. Statistical analysis was performed using the paired students *t*-test. **** p<0.0001. **E,** Hematoxylin and eosin (H&E) images of preinvasive squamous lung carcinoma (LUSC) lesions demonstrating that not all progress to invasive cancer, up to 30% regress back to normal epithelium. **F,** *TLR2* gene expression was compared between lesions of equal grade that subsequently progressed to cancer (Prog.) or regressed to normal epithelium (Reg.). Statistical analysis was performed using a linear mixed effects model to account for multiple samples from the same patient. **G,** Representative IHC staining for TLR2 in preinvasive LUSC lesions that either progressed to cancer or regressed to normal epithelium.

### *Tlr2* impairs early tumor development in murine models of lung cancer

To further characterize a possible tumor suppressor function for *TLR2* in NSCLC, we used a GEMM of preinvasive lung cancer driven by Cre-recombinase mediated activation of oncogenic *Kras^G12D^* (*Kras^LSL-G12D/+^*) (27). Following intranasal administration of Cre-recombinase expressing adenovirus (AdenoCre) lung epithelium specific activation of *Kras^G12D^* occurs and tumor formation is initiated (**Supp figure 2A**). With no alteration in the function of tumor suppressors such *Trp53*, these tumors remain low grade, consisting mainly of hyperplastic lesions, adenomas and few adenocarcinomas (grade 1-3 respectively, **Supp figure 2B**). Kras^LSL-G12D/+^ mice on either a *Tlr2* wild-type (WT) or *Tlr2* null (*Tlr2^−/−^*) background were inoculated with AdenoCre, and lung tissue was harvested between 8 and 12 weeks later for histological analysis. Kras^LSL-G12D/+^ mice on a *Tlr2^−/−^* background (*Kras^LSL-G12D/+^;Tlr2^−/−^*) developed more tumors and had a significantly increased tumor burden when compared with controls (*Kras^LSL-G12D/+^;Tlr2^+/+^*) (**Figure 2A-B**, **Supp figure 2C**). While there was no significant difference in histological grade between *Tlr2^−/−^* and control tumors (**Supp figure 2D**), *Kras^LSL-G12D/+^;Tlr2^−/−^* mice had a significant survival disadvantage in comparison to control mice (**Figure 2C**). Taken together, these data support the hypothesis that *Tlr2* loss accelerates lung tumor progression.

**Figure 2:**
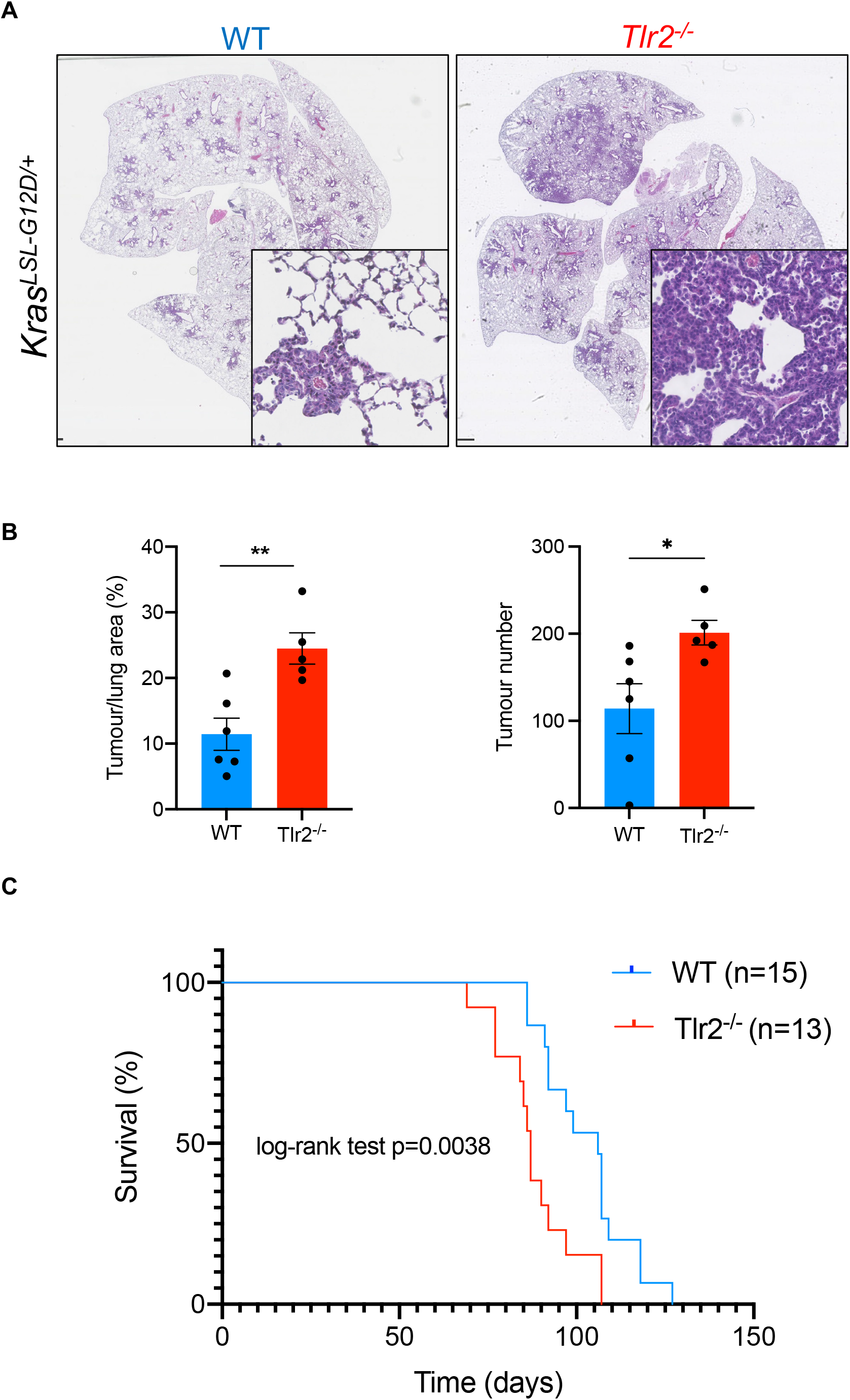
*Tlr2* has a tumor suppressor function in murine models of lung cancer. **A,** Representative hematoxylin and eosin (H&E) images from *Kras^LSL-G12D/+^* mice on either a wild-type (WT) or *Tlr2* null (*Tlr2^−/−^*) background 12-weeks following intranasal inoculation with AdenoCre. **B,** Quantification of tumor number and tumor burden (tumor area/total lung area × 100) from mice described in **A**. n=5-6 mice per group. Statistical analysis was performed using the students *t*-test. ns – non-significant, *p<0.05, **p<0.01. **C,** Kaplan-Meier curve showing survival analysis of WT or *Tlr2^−/−^ Kras^LSL-G12D/+^* mice after inoculation with AdenoCre. Statistical analysis was performed using the log-rank (Mantel-Cox) test.

### *Tlr2* controls cell intrinsic tumor suppressor responses in lung cancer initiation

We previously showed that *TLR2* regulates OIS (22) therefore we wanted to determine whether *Tlr2* loss impairs OIS in GEMM tumors. A key feature of OIS is proliferative arrest (28), therefore we analysed proliferation in lung tumors using IHC staining for the proliferation marker Ki67. Strikingly, *Tlr2^−/−^* tumors had significantly increased expression of Ki67 in comparison to control tumors (**Figure 3A**). To determine whether the increased proliferation caused by *Tlr2* loss was indeed due to impaired OIS we analysed the expression of markers of the key senescence associated events such as p53 induction and *Cdkn2a* locus activation (12,29). The cell cycle inhibitor p21 (product of *Cdkn1a* and indicator of p53 transcriptional activity) and Arf (the alternate reading frame of the *Cdkn2a* locus) were analysed in lung tumors by IHC. *Tlr2^−/−^* tumors exhibited significantly reduced expression of p21 in comparison to controls (**Figure 3A**), confirming that *Tlr2* loss impairs OIS. However, there was no significant difference in *Cdkn2a* expression (**Figure 3A**) suggesting that direct p53-p21 signalling is central to *Tlr2* mediated proliferative arrest in this context. We then wanted to assess whether the overall tumor suppressor activities of *Tlr2* were entirely dependent on p53-p21 signalling. To do this we used a GEMM whereby mice not only possess the conditional *Kras^G12D^* allele, but also have loxP sites flanking the *Trp53* allele (*Kras^LSL-G12D/+^;Trp53^fl/fl^* – so called ‘KP’ mice (27)). Therefore, intranasal administration of AdenoCre in KP mice not only activates oncogenic *Kras^G12D^* signalling but also deletes *Trp53* at the lung epithelium (**Supp figure 3A**). Unexpectedly, lung tumor burden, but not the number of tumors, was also significantly increased in KP mice on a *Tlr2^−/−^* background (*Kras^LSL-G12D/+^;Trp53^fl/fl^;Tlr2^−/−^*) (**Figure 3C-D, Supp figure 3B**), indicating that *Tlr2* has a negative effect on tumor growth in the absence of p53. Furthermore, *Tlr2* loss increased cell proliferation in KP lung tumors as determined by Ki67 immunostaining (**Figure 3C-D**) and resulted in tumors of a significantly more advanced histological grade in comparison to controls (**Figure 3E**). Taken together, these data suggest that the tumor suppressor effects of *Tlr2* extend beyond cell autonomous regulation of p53-p21 signalling.

**Figure 3:**
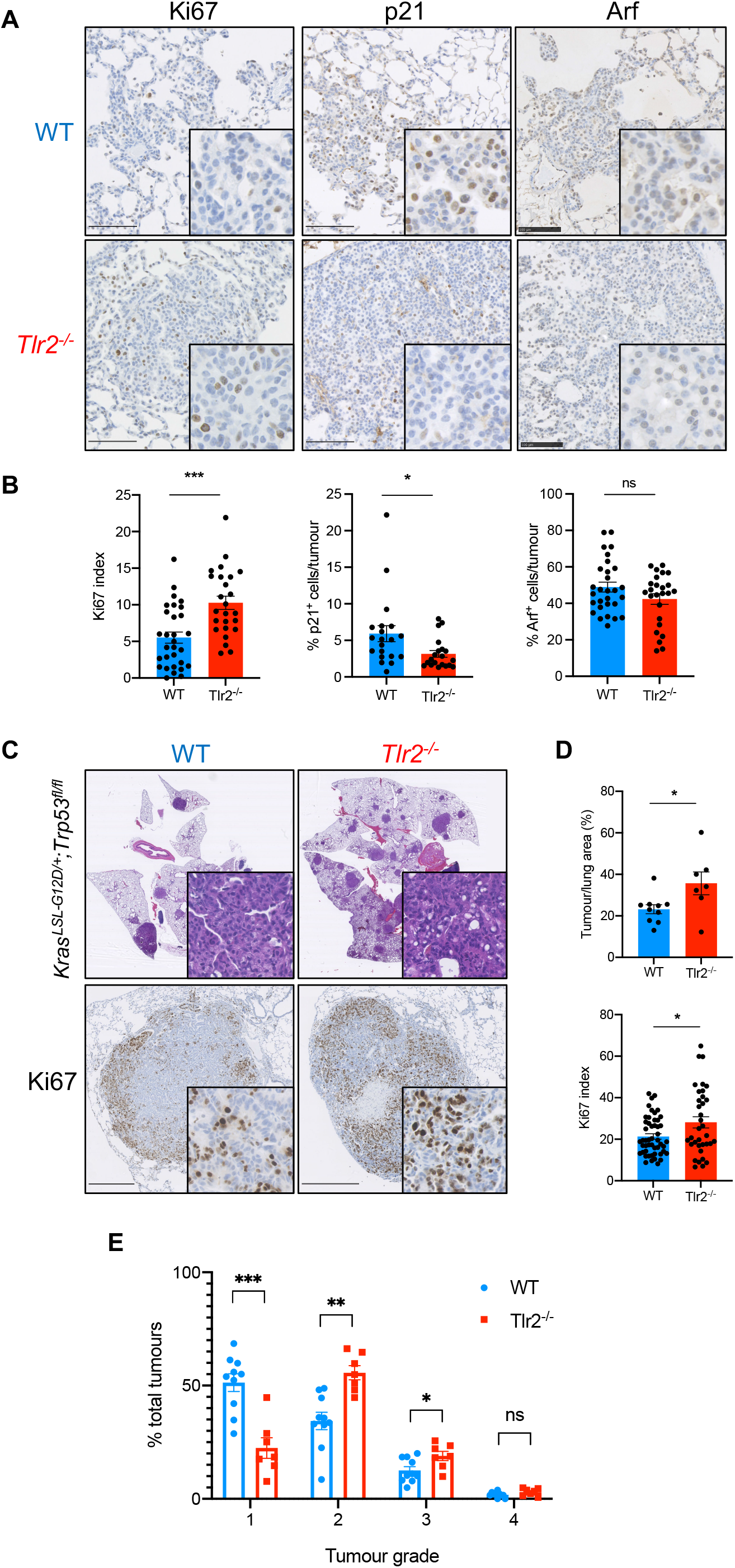
*Tlr2* loss increases proliferation and impairs senescence in lung tumors. **A,** Representative immunohistochemistry (IHC) staining of lung tumors from WT or *Tlr2^−/−^ Kras^LSL-G12D/+^* mice for Ki67, p21 and Arf, with corresponding quantification in **B**. n=5-6 mice per group (five tumors per mouse analyzed). **C,** Representative H&E and Ki67 IHC staining of lung tumors from *Kras^LSL-G12D/+^;Trp53^fl/fl^* (KP) mice on either a wild-type (WT) or *Tlr2* null (*Tlr2^−/−^*) background, with corresponding quantification in **D**. n=7-10 mice per group (five tumors per mouse analyzed). **E,** Histological grading (1–4) of tumors from *Kras^LSL-G12D/+^;Trp53^fl/fl^* mice on either a wild-type (WT) or *Tlr2* null (*Tlr2^−/−^*) background 12-weeks following intranasal inoculation with AdenoCre. n= 7-10 mice per group. Statistical analysis was performed using the students *t*-test. ns – non-significant. *p<0.05, **p<0.01, ***p<0.001.

### Epithelial cell *Tlr2* expression controls the inflammatory response during NSCLC initiation

A key non-cell autonomous facet of OIS that contributes to tumor suppression is expression of the SASP (13,17,19,20). We therefore assessed whether *Tlr2* loss impairs expression of key SASP factors including IL-1*α*, IL-1*β* and the TLR2 damage-associated molecular pattern (DAMP) A-SAA using IHC staining. We found that *Tlr2^−/−^* lung tumors had significantly reduced expression of all three SASP factors (**Figure 4A**). Furthermore, when we repeated this staining in KP lung tumors we again saw significantly reduced expression in *Tlr2^−/−^* samples (**Figure 4B**), confirming that *Tlr2* activates proinflammatory signalling independently of *Trp53*. Given the key role Tlr2 plays in innate immune sensing, we wanted to determine whether the effect of *Tlr2* loss was intrinsic to epithelial cells or due to global (principally immune cell) *Tlr2* loss. To do this we performed *in vivo* somatic genome editing using CRISPR/Cas9 to assess the effect of *Tlr2* deletion in epithelial cells only. We used the pSECC lentivirus system (30) which permits co-expression of Cre-recombinase and CRISPR/Cas9 machinery, allowing deletion of *Tlr2* and concurrent activation of oncogenic *Kras^G12D^* signalling in lung epithelial cells alone (**Supp Figure 4A**). The efficacy of our *Tlr2* guide RNA (gTlr2) expressing pSECC virus was confirmed by infecting mouse embryonic fibroblasts (MEFs) with either a control (gTomato) or gTlr2-pSECC lentivirus and measuring *Tlr2* RNA and protein expression via qRT-PCR and western blot respectively (**Supp figure 4B-C**). *Kras^LSL-G12D/+^* mice were inoculated with either gTomato or gTlr2 expressing pSECC lentivirus and lung tissue was harvested for IHC analysis as described above. We saw increased expression of the proliferation marker Ki67 in *Tlr2* targeted tumors (**Figure 4C-D**), confirming its cell autonomous role in proliferation. In concert with this, we saw that tumors from mice that received the gTlr2-pSECC lentivirus expressed significantly lower levels of SASP factors than tumors from mice that received gTom-pSECC lentivirus (**Figure 4C-D**) confirming that epithelial *Tlr2* regulates the expression of SASP. To investigate whether *TLR2* regulates the SASP in human lung cancer we performed IHC staining for TLR2 and the key SASP factor IL1*β* on consecutive sections from human LUAD samples. Not only did we observe significantly increased IL1*β* expression in LUAD epithelium compared to normal epithelium (**Supp figure 5A**), but we observed striking overlap of TLR2 and IL-1*β* expression (**Figure 5A**). We then performed automated H-score analysis for TLR2 and IL-1*β* on identical tumor regions from consecutively stained sections and found significant positive correlation between TLR2 and IL-1*β* expression (**Figure 5B**), suggesting that TLR2 can regulate the expression of the SASP in LUAD epithelium. We again analysed RNA sequencing data from preinvasive (AIS/MIA) and invasive LUAD samples (24) and found that preinvasive lesions express higher levels of the key SASP factors IL1A, IL1B and IL6 (**Supp figure 5B-D**), suggesting that TLR2-SASP signalling may oppose LUAD progression. We then analysed the expression of the *TLR2* regulated SASP and activation of the acute phase response (APR) (which includes A-SAA and was identified in our previous work (22)) in longitudinally tracked preinvasive LUSC lesions. Strikingly we found that expression of these SASP and APR factors are significantly increased in lesions that subsequently regress to normal epithelium (**Figure 5C-D and Supp figure 6A-B**). Furthermore, expression of this SASP and the APR significantly correlated with *TLR2* expression (**Figure 5E and Supp figure 6C**). We then performed IHC staining for TLR2 and IL-1*β* on serial sections from preinvasive LUSC samples, and again observed that not only is IL-1*β* expression increased in regressive lesions compared to progressive lesions (**Supp figure 6D**) but this expression is highly epithelial in origin and markedly overlaps with TLR2 expression (**Figure 5F**). Furthermore, automated H-score analysis on consecutive sections revealed significant positive correlation between epithelial TLR2 and IL-1*β* expression in preinvasive LUSC samples **(Figure 5G)**, suggesting that *TLR2* regulates expression of the SASP, aiding clinical regression of preinvasive LUSC lesions. Interestingly, *TP53* mutations typically occur early in preinvasive LUSC lesions (26), further supporting our GEMM data that demonstrates that the *TLR2* regulated SASP functions independently of *TP53*. Indeed, we saw no significant correlation between *TLR2* and *TP53* expression in the preinvasive LUSC dataset, suggesting that these entities signal independently in this context (**Supp figure 6E**). Taken together, these data suggest that TLR2 regulates the SASP in human NSCLC correlating with the impairment of tumor progression.

**Figure 4:**
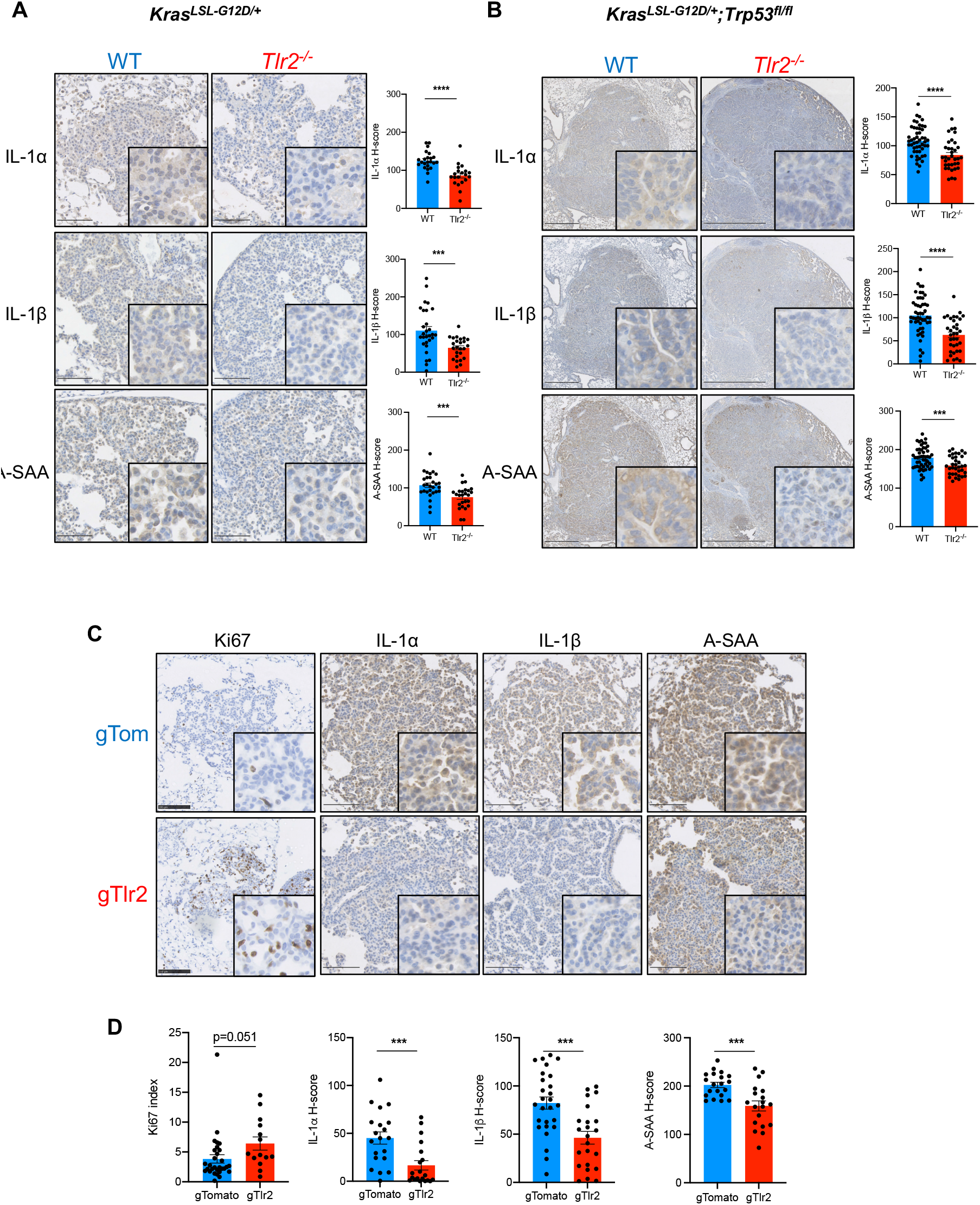
*Tlr2* loss impairs expression of the senescence associated secretory phenotype (SASP) in lung tumors. **A,** Representative IHC staining of lung tumors from *Kras^LSL-G12D/+^* mice on either a wild-type (WT) or *Tlr2* null (*Tlr2^−/−^*) background for the SASP factors interleukin-1-alpha (IL-1*α*), interleukin-1-beta (IL-1*β*) and acute-phase serum amyloid A (A-SAA), with corresponding quantification. n=5-6 mice per group (five tumors per mouse analyzed). **B,** Representative IHC staining of lung tumors from *Kras^LSL-G12D/+^;Trp53^fl/fl^* mice on either a wild-type (WT) or *Tlr2* null (*Tlr2^−/−^*) background for the same SASP factors as described in **A**, with corresponding quantification. n=7-10 mice per group (five tumors per mouse analyzed). **C,** Representative IHC staining for Ki67 and the SASP factors interleukin-1-alpha (IL-1*α*), interleukin-1-beta (IL-1*β*), acute-phase serum amyloid A (A-SAA) on lung tumors from Kras^LSL-G12D/+^ mice that either received a non-target pSECC lentivirus (gTom) or *Tlr2* targeting pSECC lentivirus (gTlr2), with corresponding quantification in **D.** n=8 mice per group. Statistical analysis was performed using the students *t*-test. ***p<0.001, ****p<0.0001.

**Figure 5:**
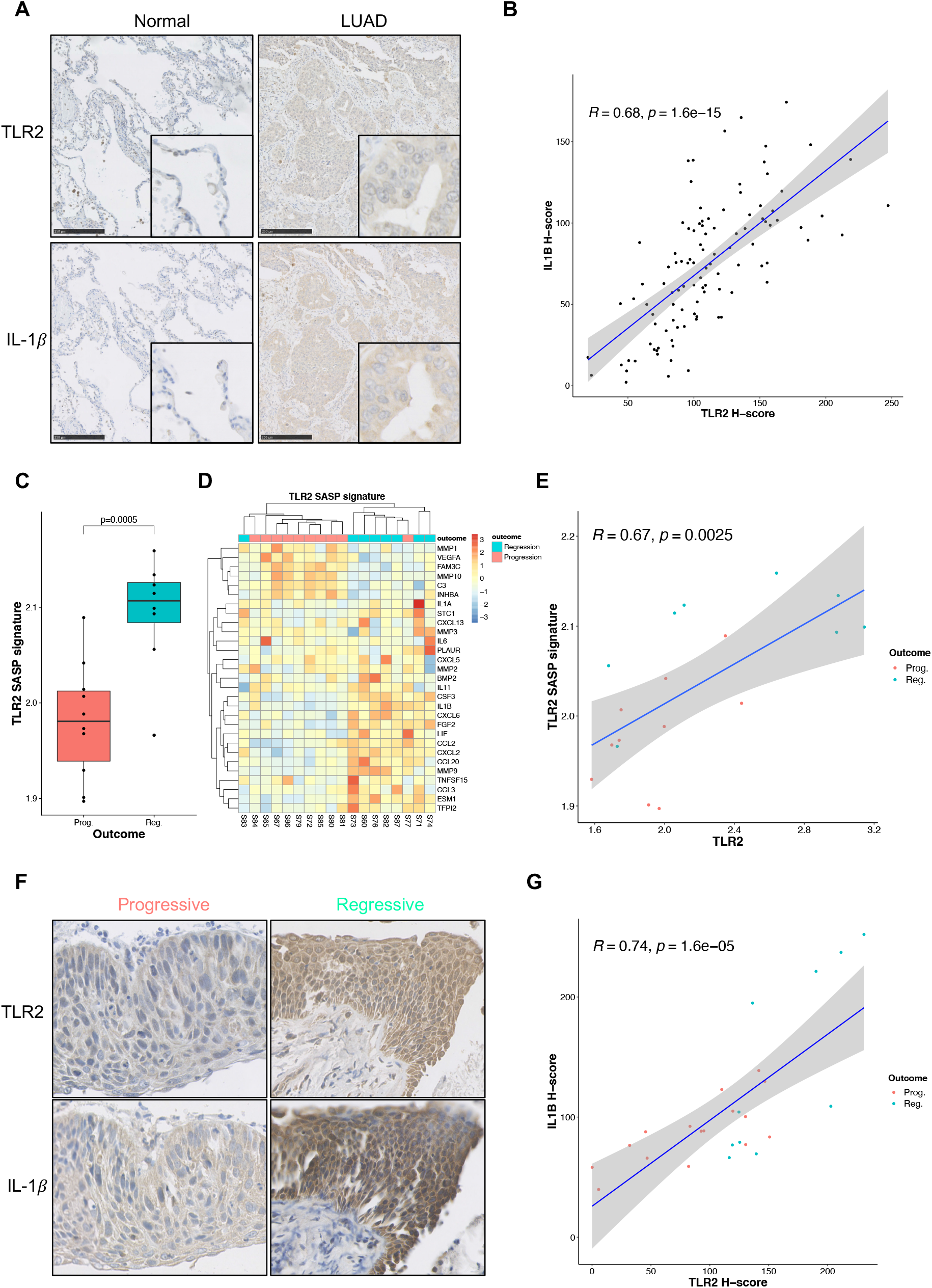
SASP expression is correlated with TLR2 expression and is associated with clinical regression in human preinvasive lung cancer. **A,** Representative IHC staining for TLR2 and IL-1*β* in consecutive sections of human LUAD and paired normal tissue, revealing distinctly overlapping epithelial expression of TLR2 and IL-1*β*. **B,** Scatter plot with Pearson correlation analysis of IHC H-score analysis performed on serial sections for TLR2 and IL-1*β*. **C,** The TLR2-SASP signature was compared between lesions of equal grade that subsequently progressed to cancer (Prog.) or regressed to normal epithelium (Reg.). Statistical analysis was performed using a linear mixed effects model to account for samples from the same patient. **D,** Heatmap demonstrating TLR2-SASP gene expression with clear clustering of progressive and regressive lesions. **E,** Scatter plot with Pearson correlation analysis comparing gene expression of *TLR2* and the TLR2-SASP signature in progressive (Prog.) and regressive (Reg.) lesions. **F,** Representative IHC staining for TLR2 and IL-1*β* in preinvasive LUSC lesions that either progressed to cancer or regressed to normal epithelium. **G,** Scatter plot with Pearson correlation analysis from IHC H-score analysis performed on serial sections for TLR2 and IL-1*β* on preinvasive LUSC lesions that either progressed to cancer (Prog.) or regressed to normal epithelium (Reg.).

### Tlr2-SASP promotes tumor immune cell recruitment and senescence surveillance in lung cancer initiation

Contributing to the tumor suppressor effect of the SASP is the recruitment of immune cells (predominantly those of the monocyte/macrophage lineage) that have been shown to clear senescence cells (so called ‘senescence surveillance’) (18,19). Therefore, we wanted to assess whether impairment of Tlr2-SASP expression impaired recruitment of immune cells to lung tumors. To do this we performed flow cytometry analysis on single cell suspensions from tumor bearing lungs from control and *Tlr2^−/−^ Kras^LSL-G12D/+^* mice (31) (**Supp figure 7A**). We saw a significant reduction in total immune cells (CD45^+^) in *Tlr2^−/−^* tumor bearing lungs (**Figure 6B**), and this was primarily driven by a reduction in alveolar macrophages (SiglecF^+^CD11c^+^) (**Figure 6A-B**). Furthermore, reduced tumor infiltration of monocyte/macrophages was confirmed in *Tlr2^−/−^* lung tumors using IHC staining for the monocyte/macrophage marker CD68 (**Figure 6C**). Tumors initiated by *Tlr2* targeting pSECC lentiviruses also exhibited significantly lower infiltration of CD68^+^ cells in comparison to non-target controls (**Figure 6D**), and local delivery of recombinant SASP factors (rIL-1*α* and rIL-1*β*) into the lungs of *Tlr2^−/−^* mice induced a robust increase in CD68^+^ cells in the lung compared to control (**Figure 6E**), confirming that *Tlr2^−/−^* monocyte/macrophages are capable of responding to local SASP factors and epithelial Tlr2-SASP expression controls the recruitment of monocyte/macrophages to lung tumors. We did not observe a significant difference in lymphoid cell recruitment in *Tlr2^−/−^* tumor bearing lungs via flow cytometry analysis (**Supp figure 7B-C**), and this was confirmed using IHC staining for the T-cell markers CD3, CD4 and CD8 (**Supp figure 7D**). Our flow cytometry experiments included analysis of T cells, a population of lung resident innate lymphoid cells that have been shown to promote lung tumorigenesis in response to altered lung commensal microbiota (32). We observed no significant increase in T cells in *Tlr2^−/−^* tumor bearing lungs suggesting that the tumor promoting effect of *Tlr2* loss is not mediated via T cell expansion (**Supp figure 7B-C**). To further determine the functional relevance of the impaired immune response following *Tlr2* loss, we took advantage of a well-characterised model of senescence surveillance, whereby oncogenes are expressed in murine hepatocytes via hydrodynamic tail vein delivery of transposable elements (19). Hydrodynamic delivery of an oncogenic *Nras* expressing plasmid (*Nras^G12V^*-GFP) was performed in WT and *Tlr2^−/−^* mice. An effector loop mutant incapable of downstream oncogenic signalling (*Nras^G12V/D38A^*-GFP) served as a negative control. 6 days after delivery, expression of the active oncogene construct resulted in a robust immune infiltrate consisting mainly of monocyte/macrophages (F4/80^+^ cells) that surrounded *Nras^G12V^* expressing hepatocytes, however this was markedly impaired in *Tlr2^−/−^* mice (**Supp figure 8A**). This impaired response corresponded with a significant impairment in clearance of senescent *Nras^G12V^* expressing hepatocytes at 12 days (**Figure 6F**). To further confirm this result, we cloned a luciferase expressing reporter construct into the *Nras^G12V^* expressing plasmid in place of GFP (*Nras^G12V^*-Luc). This approach allowed longitudinal monitoring of *Nras^G12V^* expressing cells via *in vivo* bioluminescence imaging and demonstrated a similar impairment in immune mediated clearance in *Tlr2^−/−^* mice (**Supp figure 8B-C**). Altogether, these results suggest that Tlr2 controls senescence immune surveillance during lung cancer initiation.

**Figure 6:**
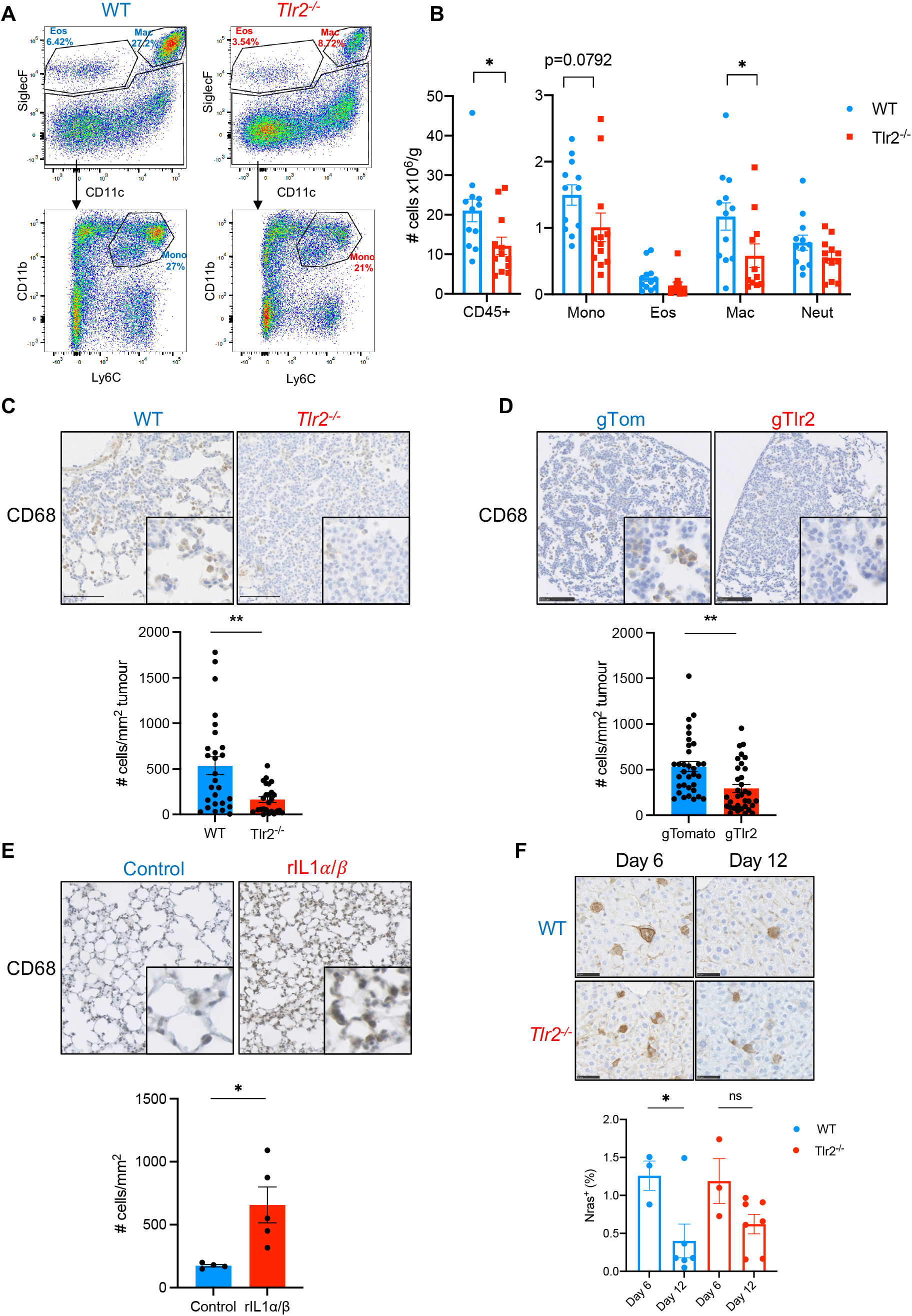
*Tlr2* loss impairs tumor immune cell recruitment and senescence surveillance. **A,** Representative flow cytometry analysis plots of myeloid populations from whole lung single cell suspensions from tumor bearing WT or *Tlr2^−/−^* mice. Percentage denotes percentage of parent population. **B,** Corresponding quantification of total immune cells (CD45^+^) and myeloid cells from WT (blue) and *Tlr2^−/−^* (red) mice. Mono – monocytes, Eos – eosinophils, Mac – Macrophages, Neut – Neutrophils. n=12 mice per group. **C,** Representative IHC staining for the monocyte/macrophage marker CD68 in lung tumors from WT or *Tlr2^−/−^* mice with corresponding quantification. n=5-6 mice per group (five tumours per mouse analyzed). **D,** Representative IHC staining for the CD68 in lung tumors from *Kras^LSL-G12D/+^* mice inoculated with either gTomato (gTom) or gTlr2 expressing lentivirus, with corresponding quantification. **E,** Representative IHC staining for CD68 in lung tissue from *Tlr2^−/−^* mice intranasally inoculated with either PBS (control) or recombinant interleukin-1-alpha (rIL-1*α*) and recombinant interleukin-1-beta (rIL-1*β*), with corresponding quantification. **F,** Liver IHC staining for Nras from WT or *Tlr2^−/−^* mice six and twelve days after hydrodynamic delivery of *Nras^G12V^* expressing transposons, with corresponding quantification. Statistical analysis was performed using the students *t*-test. ns – non-significant, *p<0.05, **p<0.01.

### Pharmacological intervention with Tlr2 agonists prevents early lung tumor growth

Given the tumor suppressor function we identified for Tlr2-SASP signalling in the lung, we wanted to investigate whether we can inhibit lung tumor growth by pharmacologically activating Tlr2 signalling. We used the synthetic Tlr2 agonist Pam2CSK4 delivered via nebulisation to allow direct repeated administration of drug to the lung epithelium. *Kras^LSL-^ G12D/+* mice on a *Tlr2* WT background were intranasally inoculated with AdenoCre as previously described. Two-weeks later they were subjected to weekly dosing with nebulized Pam2CSK4 or vehicle control (0.9% sodium chloride) for 8 weeks, prior to sacrifice and lung tumor analysis (**Figure 7A**). While tumor number was unchanged, tumor burden was significantly reduced in mice that received Pam2CSK4 in comparison to controls (**Figure 7B**) indicating that *Tlr2* activation inhibits lung tumor growth. This was associated with significantly increased p21 expression in Pam2CSK4 treated tumors, however Ki67 expression was unchanged **(Figure 7C)**, suggesting a higher contribution for senescence immune surveillance following treatment with a Tlr2 agonist. The expression of the SASP factors IL-1*α*, IL-1*β* and A-SAA in tumors following Pam2CSK4 treatment was also increased (**Figure 7D**), indicating increased SASP expression following *Tlr2* activation. We saw no change in CD3^+^ T-cell infiltration following Pam2CSK4 treatment, however we did observe a dramatic reduction in CD68^+^ macrophage/monocyte infiltration (**Supp figure 9A**). This unexpected finding is likely explained by Toll-like receptor tolerance which is a well-recognised phenomenon in monocyte derived populations via repression of NF-κB signalling (33). We repeated this experiment in *Tlr2^−/−^* mice and saw no significant difference in tumor burden, Ki67 or SASP expression (**Supp figure 9B**), confirming that Pam2CSK4 acts via Tlr2 to mediate its tumor suppressor effect. Taken together, our data show that activation of Tlr2 can inhibit early lung tumor growth, highlighting a novel therapeutic target for the treatment of early-stage NSCLC.

**Figure 7:**
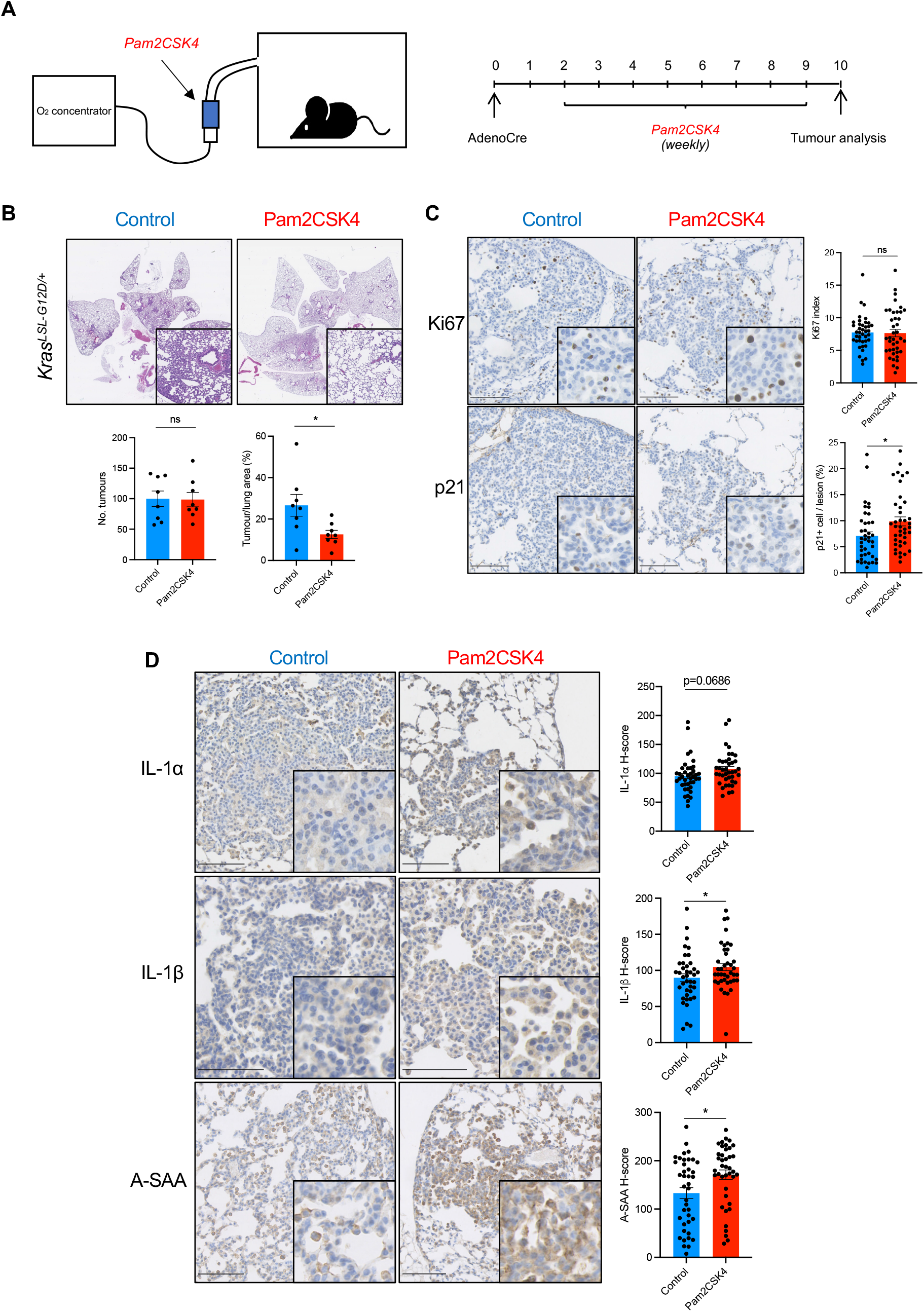
TLR2 activation inhibits lung tumor growth. **A,** Kras^LSL-G12D/+^ mice were inoculated with AdenoCre as described previously. Two weeks after inoculation they were subjected to weekly nebulized dosing with Pam2CSK4 or control (0.9% NaCl solution) for eight weeks, prior to sacrifice and histological analysis. **B,** Representative H&E staining of lung sections from Kras^LSL-G12D/+^ mice after either control or Pam2CSK4 treatment, with corresponding tumor number and tumor burden analysis. n=8 mice per group. **C,** Representative IHC staining for Ki67 and p21 in lung tumors from Kras^LSL-G12D/+^ after either control or Pam2CSK4 treatment, with corresponding quantification. n=8 mice per group (five tumors per mouse analyzed). **D,** Representative IHC staining for the SASP factors interleukin-1-alpha (IL-1*α*), interleukin-1-beta (IL-1*β*) and acute-phase serum amyloid A (A-SAA), with corresponding quantification. n=8 mice per group (five tumors per mouse analyzed). Statistical analysis was performed using the students *t*-test. *p<0.05.

## Discussion

We have shown here for the first time that TLR2 has a critical role in NSCLC initiation via regulation of both cell intrinsic tumor suppressor responses and SASP mediated immune surveillance. Using clinical samples, we have demonstrated that *TLR2* and the SASP are highly expressed in the tumor epithelium of both the main subtypes of human NSCLC (lung adenocarcinoma (LUAD) and squamous cell carcinoma (LUSC)). By analysing the transcriptome of LUAD samples we have shown that high *TLR2* expression significantly correlates with improved prognosis and expression drops during progression from the preinvasive to the invasive stages of carcinogenesis, suggesting a tumor suppressor function for *TLR2* in this context. Furthermore, epithelial TLR2 expression significantly correlates with the expression of key SASP factors, indicating a potential regulatory role for TLR2 in activating proinflammatory responses in human LUAD. Unfortunately, we were unable to accurately determine the prognostic significance of *TLR2* expression in LUAD precursor lesions as these lesions are treated by surgical resection, precluding longitudinal follow up of lesion outcome. To overcome this, we analysed a unique gene expression dataset from human preinvasive LUSC samples that had undergone longitudinal clinical and molecular follow-up, detailing either progression to invasive malignancy or histological regression (26). Strikingly, we demonstrated that the expression of *TLR2* and the SASP was significantly associated with subsequent clinical regression and *TLR2* expression positively correlated with SASP expression, further supporting the hypothesis that *TLR2* orchestrates an anti-tumor response in early NSCLC.

To confirm these observations and to glean mechanistic insight into the tumor suppressor function of TLR2, we used a *Kras^G12D^* driven GEMM of NSCLC. Using this we confirmed that *Tlr2* loss is associated with accelerated lung tumorigenesis and shortened survival. *Kras* mutations are infrequent in human LUSC (as opposed to being the most frequently mutated oncogene in LUAD). It could be argued therefore that correlation between our *Kras^G12D^* driven GEMM/LUAD and human preinvasive LUSC is not valid. However, the upregulation of *TLR2* and the SASP is not confined to oncogenic *RAS* induced senescence alone. Our previous work identified similar upregulation in multiple forms of senescence associated with genotoxic stress (including therapy-induced senescence, mutant *BRAF* induced senescence in melanoma, DNA damage induced senescence and replicative senescence) (22). Furthermore, similar epithelial specific toll-like receptor inflammatory signalling has been identified in murine prostate intraepithelial neoplasia (PIN) caused by *Pten* deletion (34) and has been shown to mediate senescence in tubular epithelial cells (TECs) in the murine kidney after injury (35).

We observed increased p21 expression in *Tlr2* wild-type GEMM lung tumors in comparison to *Tlr2* null counterparts and also observed increased p21 expression following treatment with a *Tlr2* agonist, suggesting that *Tlr2* reinforces the growth arrest via p53-p21 signalling. However, following concurrent *Kras^G12D^* activation and *Trp53* loss (using KP mice), *Tlr2* null tumors still exhibit accelerated proliferation and advanced histological grade, confirming that the tumor suppressor effect of *Tlr2* is not solely dependent on p53-p21 signalling. Indeed, we demonstrated that expression of the SASP was not dependent on *Trp53*, leading us to speculate that the non-cell autonomous tumor suppressor functions of the SASP may be underlying this effect (see below). Interestingly, suppression of p16 has recently been shown to perturb expression of the SASP following oncogene activation (36) suggesting that intact p16-Rb signalling in our *Trp53* null tumors may maintain SASP expression. It has also been shown that mutant *TRP53* interferes with cGAS-STING-TBK1 signalling, subsequently impairing the innate immune response and promoting cancer progression (37). This effect is not observed in *Trp53* null cells, suggesting that mutant *TRP53* exerts gain of function activities to inactivate tumor immune surveillance. cGAS-STING activation is a key event in expression of the SASP mediated via cytoplasmic chromatin sensing (38,39) and we demonstrated in our previous work that TLR2 functions downstream of this activation (22). Therefore, it remains to be determined whether intact cGAS-STING activation may explain the persistent effect of *Tlr2* loss we observe in our *Trp53* null tumors.

The clearance of senescent cells by immune cells is a key non-cell autonomous tumor suppressor function of the SASP (18,19). This characteristic was first recognised by Xue et al who demonstrated a robust innate immune cell mediated clearance of hepatocarcinomas upon reactivation of the senescence program (18). Further delineation of this process revealed that senescence induced by oncogene activation leads to expression of pro-inflammatory SASP factors, which orchestrate an adaptive immune response termed ‘senescence surveillance’. This response is dependent on CD4^+^ T-cells however recruited monocyte/macrophages act as the effector cells mediating clearance of senescent cells (19). Corresponding with the reduction in SASP expression in our GEMM lung tumors, we identified a significant reduction in total immune cells in tumor bearing lungs from *Kras^G12D^;Tlr2^−/−^* mice. This was accounted for by a significant reduction in alveolar macrophages and reduced recruited monocytes. While we saw a trend towards reduced T-cell recruitment/infiltration, these changes were not significant. It must be noted however that tumors from *Kras^G12D^* driven GEMMs of lung cancer have few protein-altering mutations and are therefore unlikely to express significant numbers of neoantigens (40). Heightened CD8^+^ T-cell responses can be achieved by forced expression of T-cell antigens, however this is not sustained and is characterised by a poorly functioning exhausted T-cell response (41). Therefore, due to the limitation of our GEMM we cannot exclude heightened T-cell mediated immunity via the Tlr2 regulated SASP as a driver of lesion regression as seen in human NSCLC precursors (42). Our flow cytometry analysis also did not allow determination of the activation state of immune cells nor evidence of potential exhaustion. Immune cell exhaustion has been shown to be a feature in late-stage lung tumors in this model (41,43), therefore further studies to analyse the activation state of immune cells following *Tlr2* loss are warranted.

Recently it has been shown than endothelial Tlr2 signalling supports the recruitment of immune cells to tumors via expression of proinflammatory cell adhesion molecules (44), therefore it is possible that endothelial *Tlr2* loss is a contributing factor to the reduced immune infiltration we see in our global *Tlr2^−/−^* mice. However, the same research study demonstrated extensive epithelial expression of *Tlr2* in the murine lung using the TLR2-IRES-EGFP reporter mouse, suggesting that epithelial *Tlr2* could also be vital in regulating this process and explaining why we see such a specific tumor suppressor effect for *Tlr2* in lung cancer. Furthermore, we confirmed that epithelial *Tlr2* loss underpins our observed phenotype using *in vivo* somatic genome editing, allowing concurrent *Tlr2* deletion and *Kras^G12D^* activation in lung epithelial cells only.

Gene expression profiling of human preinvasive LUSC lesions has revealed an abundance of immune sensing during the early stages of tumorigenesis with activated T-cells and myeloid cells (including macrophages) peaking immediately prior to invasion (45). Immune escape mechanisms are subsequently activated, promoting the progression to invasive cancer (45). We have recently shown that immune surveillance is strongly implicated in lesion regression (42), leading us to suggests that this may be supported by the *TLR2* regulated SASP. Macrophages make up the majority of the tumor immune infiltrate in human lung cancer (46) and studies examining the microanatomical location of macrophages have revealed that high macrophage infiltration within lung tumor epithelium (vs tumor stroma) is significantly associated with improved patient survival (47–49). It is therefore tempting to speculate that activated TLR2-SASP signalling in lung tumors instruct the recruitment of macrophages to the epithelium to subsequently impair tumor progression via immune surveillance.

Lastly, to determine whether our findings have therapeutic potential, we tested whether activation of Tlr2 could perturb lung tumor growth. We found that following inhalational delivery of a synthetic Tlr2 agonist tumor growth was significantly reduced and this corresponded with an increased expression of p21 and key SASP factors. We have identified for the first time a tumor suppressor role for *TLR2* in NSCLC, via orchestrating both cell intrinsic tumor suppression and SASP induced immune surveillance. Pharmacological modulation of this pathway may provide a novel therapeutic approach for the treatment of early-stage lung cancer. *TLR2* and the associated SASP are expressed by preinvasive tumor lesions. Given that these factors are readily secreted, they are likely to represent candidate biomarkers of preinvasive disease measured either directly from patient plasma or more specifically via tumor/organ specific extracellular vesicles (50). This approach could be used to identify and target lung cancer earlier, or perhaps stratify lung cancer screening populations.

## Methods

### Animal experiments

All mice were housed in a specific pathogen free environment with food and water *ad libitum* in accordance with UK home office guidance at the Biomedical Research Facility, University of Edinburgh. Black six (C57BL/6) mice harbouring the conditional oncogenic *Kras^G12D^* allele (Kras^G12D/+^) were purchased from the Jackson Laboratory (jax.org) and interbred with black six (C57BL/6) mice lacking *Tlr2* (*Tlr2^−/−^*) (received from Dr Jen Morton, University of Glasgow) to generate our *Kras^G12D/+^;Tlr2^−/−^* line. This line was then interbred with black six (C57BL/6) mice harbouring the ‘floxed’ *Trp53* allele (*Trp53^fl/fl^*) (received from Dr Luke Boulter, University of Edinburgh) to generate our *Kras^G12D/+^;Trp53^fl/fl^;Tlr2^−/−^* line. Mice between the ages of 8-12 weeks were anaesthetised with medetomidine and ketamine prior to intranasal inoculation with 40ul of virus solution (1.5×10^7 plaque forming units (PFU) of adenovirus expressing Cre-recombinase under control of the CMV promoter suspended in minimum essential media (MEM), or pSECC lentivirus (see below)). Mice were humanely sacrificed 8-15 weeks later. Lung tissue was harvested, inflated with 10% neutral buffered formalin (NBF) and fixed in NBF overnight prior to tissue processing and paraffin embedding. For recombinant IL1α/IL1β inoculation, mice were anaesthetised as described above and intranasally inoculated with 100ng of carrier free recombinant protein (rIL-1*α* – Biolegend #575002, rIL-1*β* – Biolegend #575102) suspended in sterile phosphate buffered saline (PBS). For Tlr2 agonist experiments, 100ug Pam2CSK4 was diluted in 3mls 0.9% NaCl and delivered via a nebuliser to mice housed in a nebuliser chamber over 30 minutes. Control mice received 0.9% NaCl only. This was performed weekly for eight weeks prior to humane sacrifice and tissue processing as above. For hydrodynamic tail vein injection experiments, wild-type (WT) and Tlr2 null (*Tlr2^−/−^*) mice on a C57BL/6 background aged between 6-12 weeks were included in the study. DNA plasmids were prepared using the Qiagen Plasmid Maxi Kit (Qiagen, Germany) as per the manufacturer’s instructions. Each mouse received 6μg of a sleeping beauty transposase expressing plasmid (CMV-SB13 transposase) and 20μg of transposon (pT3-Nras^G12V^-IRES-GFP/Luc or pT3-Nras^G12V/D38A^-IRES-GFP) encoding plasmid diluted in 0.9% NaCl solution to 10% of the animal’s body weight delivered via the lateral tail vein within 10 seconds. For longitudinal bioluminescence imaging mice were administered with a 15mg/ml D-luciferin solution subcutaneously at a dose of 150mg/kg (injection volume of 0.1ml/10g body weight). Following this, animals were anaesthetised with isoflurane, abdomens shaved and imaged using an IVIS Lumina S5 *in vivo* imaging system (IVIS®). Optimal signal time was determined using a kinetic curve.

### Immunohistochemistry

3μm sections were cut onto adhesion slides from NBF fixed, paraffin embedded (FFPE) samples then placed in a 60°C oven for at least two hours. Slides were de-waxed in xylene and rehydrated in ethanol of decreasing concentrations. Antigen retrieval was performed using heated sodium citrate buffer for ten minutes. Samples were blocked with hydrogen peroxide, avidin/biotin block (if using biotinylated secondary antibodies) and protein block prior to incubation with primary antibody (**Supp table 1**). Samples were washed then incubated with appropriate biotinylated secondary antibodies followed by streptavidin-HRP conjugate (VECTASTAIN ABC reagent) then revealed with 3,3’Diaminobenzidine (DAB) diluted in DAB substrate (Abcam), with the exception of CD8 staining which was performed using HRP conjugated secondary antibodies followed by revealing with DAB. Samples were counterstained with haematoxylin before dehydration with ethanol, clearing with xylene and mounting. Immunohistochemistry slides were imaged using the Hamamatsu Nanozoomer XR microscope with NDP scan v3.1 software. Tumor burden quantification was performed in a blinded fashion on haematoxylin and eosin (H&E) stained slides using NDP viewer v2 software. Immunohistochemistry quantification (positive cell detection or H-score analysis) was performed using QuPath software v0.1.2 (51). Histological grade analysis was performed in a blinded fashion using predefined criteria (27) and validated by an independent reviewer on a subset of tumours (n=120) with strong inter-rater agreement as assessed using Cohen’s kappa coefficient (weighted Kappa 0.854, unweighted kappa 0.789 (95% CI 0.698-0.880)). Co-immunofluorescence staining was performed on 3μm FFPE samples that were dewaxed and rehydrated as above. Permeabilization was performed with a 0.1% Triton X-100 solution in PBS prior to incubation with primary antibodies. Samples were then washed followed by incubation with appropriate fluorophore conjugated secondary antibodies. Samples were then washed, stained with DAPI and mounted with Vectashield®. Samples were imaged using a confocal microscope using NIS-Elements software (Nikon).

### Flow cytometry

Tumor bearing mice were humanely sacrificed ten weeks after inoculation with Cre-expressing adenovirus. The lung vasculature was perfused with up to 30mls of ice-cold PBS over 30 seconds via the right ventricle to flush the lungs of all blood. Tissue was collected and weighed, roughly diced prior to 25-minute incubation with 10mls of a digestion enzyme mix (0.8mg/ml collagenase V, 0.625mg/ml collagenase D, 1mg/ml dispase and 30μg/ml DNase) in a shaking incubator at 37°C. Digested lung tissue was filtered through a 100μm cell strainer, washed twice with RPMI (Lonza) and incubated with 3mls red cell lysis buffer (Sigma) for three minutes. After further washing samples were filtered through a 35μm cell strainer directly into 5ml (12×75) polystyrene round-bottom FACS tubes and counted using a Muse Cell Analyser (Merck Millipore). Single cell suspensions were blocked with 10% mouse serum and 1% Fc block prior to incubation with cell surface fluorophore conjugated primary antibodies (**Supp table 2**). Samples were analysed on a BD LSR Fortessa X-20 analyser using BD FACSDiva 8.0.1 software. Data were analysed using FloJo software v10.2. Gating was designed using fluorescence minus one (FMO) samples. Compensation was performed using single antibody stained OneComp eBeads (ThermoFisher). Total cell number was determined by multiplying the percentage of live fluorophore positive cells by the total number of live cells isolated per sample and this was normalised to tissue weight.

### Molecular cloning and lentivirus generation

Empty vector pSECC plasmids were obtained from Tyler Jacks via Addgene (#60820). Guide RNA (gRNA) oligos were selected from the GeCKO library (52) and ordered from Sigma. pSECC vectors were digested with BsmBI restriction enzymes and ligated with compatible annealed oligos (**Supp table 3**). Lentiviruses were produced by polyethylenimine (PEI) mediated co-transfection of 293T cells with lentiviral backbone and packaging plasmids (PAX2 and VSVG) with pSECC plasmids. Supernatant was collected at 48- and 72-hours post transfection and virus was concentrated using Lenti-X™ concentrator (Takara) and resuspended in appropriate volumes of MEM. Lentiviral titrations were performed by infecting 3TZ cells (containing the loxP-STOP-loxP *β*-Gal cassette) followed by beta-galactosidase staining. For transposon construct cloning, the pT3-Nras^G12V^-IRES-GFP plasmid were obtained directly from Scott Lowe and an MSCV-IRES-Luciferase plasmid was obtained from Scott Lowe via Addgene (#18760). Both plasmids were digested with AvrII and AgeI-HF restriction enzymes prior to ligation and transformation into NEB Stable Competent E. coli. Positive colonies were identified with a colony PCR and subsequent sequencing.

### Human lung cancer sample analysis

Anonymous FFPE human LUAD samples were obtained with informed consent and ethical approval from the NRS Lothian Bioresource, Edinburgh. FFPE human preinvasive LUSC samples were obtained with informed consent from the pathology department at University College London Hospital (UCLH) from patient enrolled in the UCLH surveillance study. LUAD and LUSC samples were stained and analysed as described above. Protein expression correlation was performed on serial sections using QuPath (51). All human gene expression data presented were re-analysed from published datasets and sample acquisition and analysis has been previously described (24,26). Briefly, RNA-sequencing data from LUAD lesions were downloaded with permission from the European Genome-Phenome Archive (EGA) and aligned to the human genome (GRCh38) using STAR (v2.6.1) (53) and read counts were quantified using Salmon (version 1.4.0). Differential gene expression analysis was performed using the DESeq2 package on R (version 4.0.3) (54). For progressive vs regressive preinvasive LUSC sample analysis, gene expression data acquired from Illumina and Affymetrix microarray platforms was used from patients enrolled in the UCLH Surveillance study. ‘Index’ biopsies were used for gene expression analysis and were defined as a preinvasive LUSC biopsy that was performed four months prior to either a diagnostic cancer biopsy (progression) or a normal/low grade lesion biopsy (regression). Data from preinvasive LUSC gene expression were analysed using R (version 4.0.3) using a linear mixed effects model to account for multiple samples per patient. The TLR2 SASP signature and acute phase response signature were determined by calculating the geometric mean of gene expression values of each of the SASP/APR factors regulated by TLR2 as described previously (22).

### Statistical analysis

Statistical analysis was performed using R software (version 4.0.3) and GraphPad Prism (version 9.0.1). Details regarding the statistical tests used can be found in the figure legends. The statistical tests used were justified as appropriate based on sample size and distribution. Students *t*-test or Mann-Whitney tests (or equivalent tests for paired analysis) were used for two-condition comparisons, with a significance cut off of p<0.05.

## Author contributions

J.C.A, S.W, M.F and F.R.M conceived the study design. F.R.M and M.M performed all lung cancer mouse experiments. L.B and F.R.M performed hydrodynamic tail vein injection experiments. A.Q and F.R.M performed molecular cloning experiments. F.R.M and E.F performed flow cytometry experiments. A.H.S analysed TCGA survival data. A.P analysed preinvasive LUSC samples from the UCLH surveillance study. P.G, A.M and F.R.M performed bioinformatic analysis of LUAD RNA sequencing data. Pathological grading was performed by F.R.M with independent validation by J.C. Human LUAD samples were provided by W.A.H.W and human LUSC samples were provided by V.H.T and S.M.J. F.R.M and J.C.A co-wrote the manuscript. J.C.A provided overall study supervision.

## Acknowledgements

We thank all of the patients who kindly gifted clinical samples to the NRS Lothian Bioresource and took part in the UCLH surveillance study. We thank A. Finch, M. Christophorou, A. von Kriegheim, N. Gammoh and the ECAT fellowship directors for helpful criticism, discussion and encouragement. We thank the CRUK Edinburgh Centre, the Western General BRF facility staff and Histology service at the CRUK Edinburgh Centre. We thank J. Morton for the *Tlr2*^−/−^ mice and S.W Lowe for the pT3-NRas^G12V^-IRES-GFP, pT3-NRas^G12V/D38A^-IRES-GFP and CMV-SN13 plasmids. F.R.M is funded by a Wellcome Trust Clinical Research Fellowship through the Edinburgh Clinical Academic Track (ECAT) programme (203913/Z/16/Z), a Wellcome Trust-ISSF3 award (IS3-R1.07 20/21) and a Wellcome Trust iTPA award (209710/Z/17/Z). J.C.A core lab funding was received from Cancer Research UK (C47559/A16243, Training and Career Development Board – Career Development Fellowship) and the University of Edinburgh (Chancellor’s Fellowship). S.W is supported by a Cancer Research UK Senior Fellowship (A29576). J.C is supported by a Wellcome Trust Clinical Lectureship through the ECAT programme (203913/Z/16/Z).

