## Supplementary information for "Toll-like receptor 2 orchestrates a potent anti-tumor response in non-small cell lung cancer"

### Supplementary figures

#### Supplementary figure 1

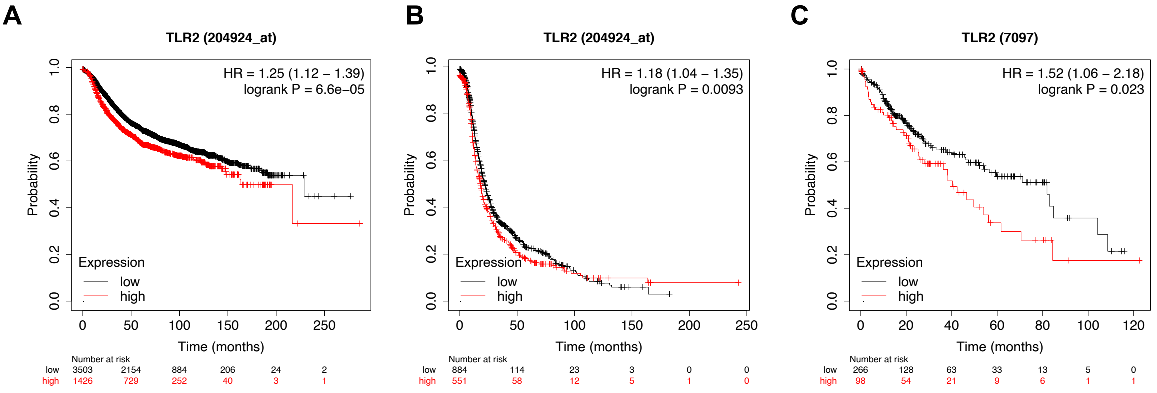


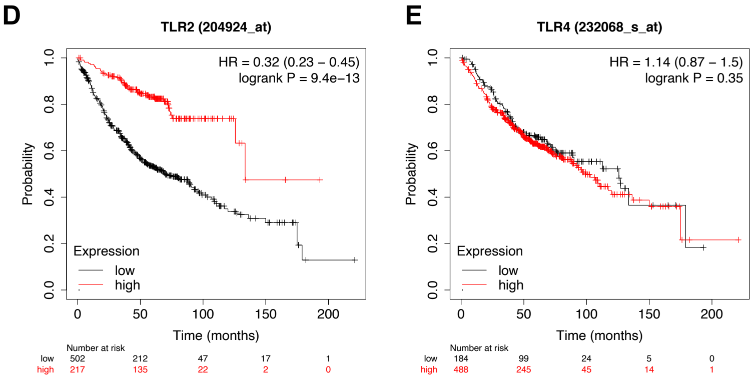


F

| Gene | HR | p-value |
|---|---|---|
| TLR1 | 0.65 | 0.0002 |
| TLR2 | 0.32 | 9.4E-13 |
| TLR6 | 0.65 | 0.0005 |
| TLR10 | 0.72 | 0.0138 |
| TLR4 | 1.14 | 0.3586 |
| RB1 | 0.3 | 7.30E-15 |

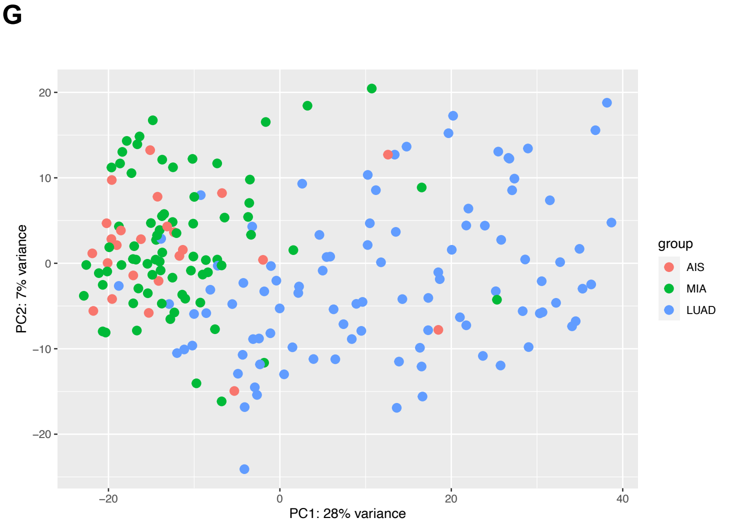


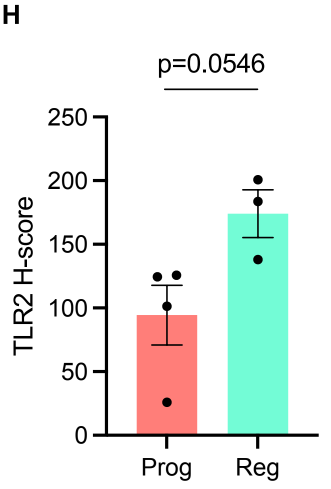

**Supplementary figure 1:** Kaplan-Meier survival plots derived from KMplot.com analyzing *TLR2* expression in **A**, Breast, **B,** ovarian and **C,** liver cancer showing no correlation between *TLR2* expression and improved survival. **D,** Kaplan-Meier survival plot showing the overall survival for *TLR2* and **E,** *TLR4* expression in lung adenocarcinoma samples from KMplot.com (corresponding to 866 patients from the GEO, EGA and TCGA). **F,** Table showing hazard ratios (HR) and corresponding p-value of each indicated gene (including *TLR2* and its dimerization partners *TLR1*, *6* and *10*, and the non-TLR2 associated plasma membrane Toll-like receptor *TLR4*) in human lung adenocarcinoma. *TLR2* shows the lowest HR, which compares with the well characterized tumor suppressor gene *RB1*. The effect in lung adenocarcinoma is specific to the TLR2 network as the other plasma membrane TLR (*TLR4*) does not show prognostic value. **G,** Principal component analysis (PCA) plot of gene expression from AIS (adenocarcinoma in situ), MIA (minimally invasive adenocarcinoma) and invasive lung adenocarcinoma (LUAD) revealing clustering of preinvasive lesions (AIS and MIA). **H,** IHC quantification of TLR2 expression in preinvasive LUSC lesions that either progressed to cancer (Prog) or regressed to normal epithelium (Reg). Statistical analysis was performed using the students *t*-test.

#### Supplementary figure 2

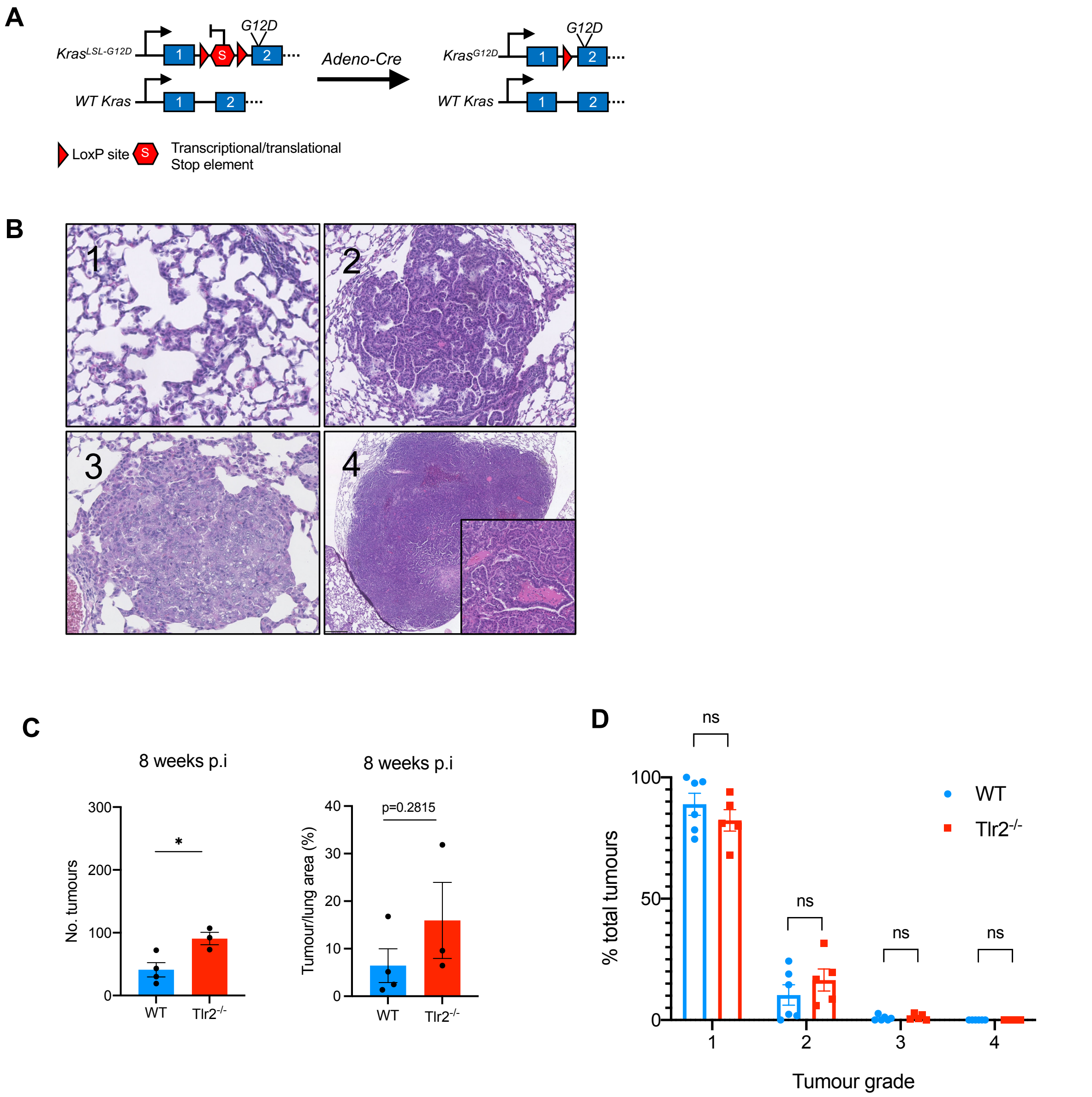

**Supplementary figure 2: A,** The *Kras*$^{LSL\text{-}G12D}$ allele consists of a STOP codon flanked by loxP sites upstream of the oncogenic *Kras*$^{G12D}$ allele. Following exposure to adenovirus expressing Cre-recombinase (AdenoCre), the STOP codon is removed resulting in constitutive activation of mutant *Kras*$^{G12D}$ signaling. **B,** Following *Kras*$^{G12D}$ activation +/- *Trp53* loss at the lung epithelium, lung tumors or varying grades develop (1 – Atypical adenomatous hyperplasia, 2 – Adenomas, 3 – Adenocarcinoma, 4 – Invasive adenocarcinoma). **C,** Quantification of tumor number and tumor burden (tumor area/total lung area x 100) from Kras$^{LSL\text{-}G12D/+}$ mice on either a wild-type (WT) or *Tlr2* null (*Tlr2*$^{-/-}$) background 8-weeks following intranasal inoculation with AdenoCre. p.i – post inoculation. n=3-4 mice per group. **D,** Histological grading (1-4) of tumors from *Kras*$^{LSL\text{-}G12D/+}$ mice on either a wild-type (WT) or *Tlr2* null (*Tlr2*$^{-/-}$) background 12-weeks following intranasal inoculation with AdenoCre. n= 5-6 mice per group. Statistical analysis was performed using the students *t*-test. ns – non-significant, *p<0.05.

#### Supplementary figure 3

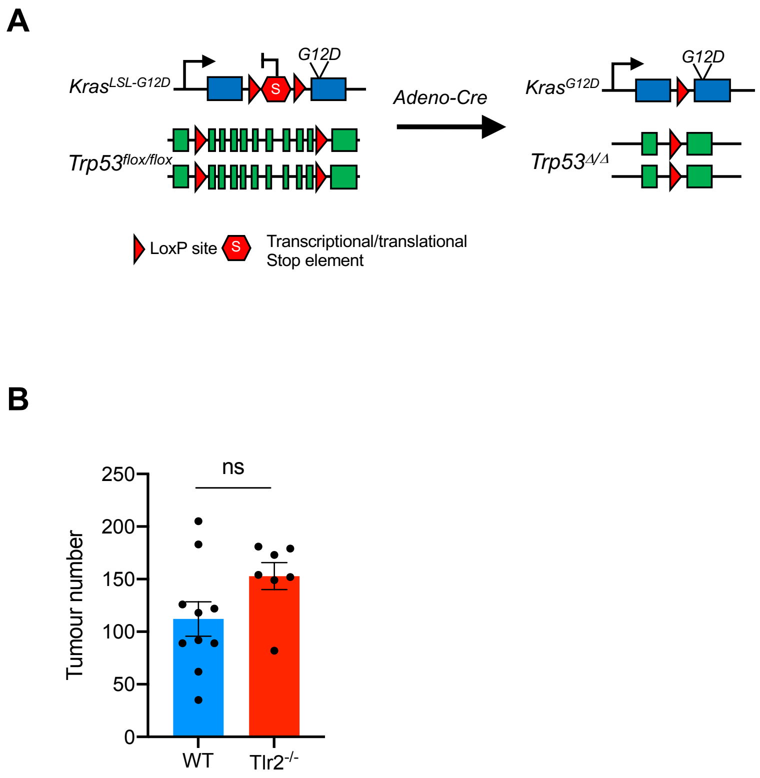


**Supplementary figure 3: A,** *Kras*$^{LSL\text{-}G12D/+}$;*Trp53*$^{fl/fl}$ mice not only harbor the *Kras*$^{LSL\text{-}G12D}$ allele but also loxP sites flanking the *Trp53* allele. Following exposure to adenovirus expressing Cre-recombinase (AdenoCre), the STOP codon is removed resulting in constitutive activation of mutant *Kras*$^{G12D}$ signaling and loss of *Trp53*. **B,** Quantification of tumor number in *Kras*$^{LSL\text{-}G12D/+}$;*Trp53*$^{fl/fl}$ mice 12 weeks after inoculation with AdenoCre. n=7-10 mice per group. Statistical analysis was performed using the students *t*-test. ns – non-significant.

Supplementary figure 4

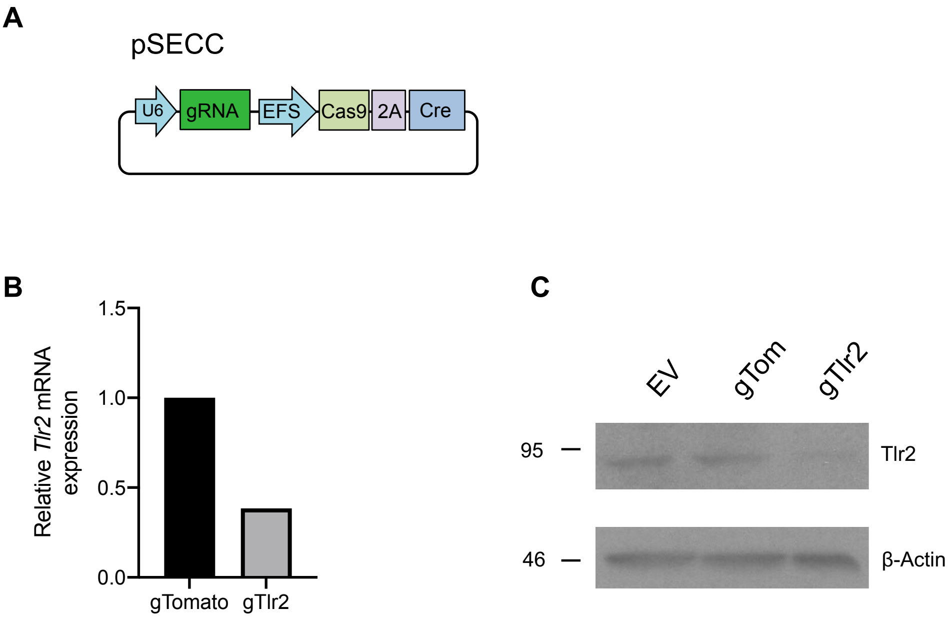


**Supplementary figure 4: A,** *Tlr2* targeting gRNA were cloned into pSECC lentiviral plasmids to allow concurrent *Tlr2* deletion and *Kras*$^{G12D}$ activation in lung epithelial cells only following intranasal inoculation of *Kras*$^{LSL\text{-}G12D/+}$ mice. **B,** qRT-PCR and **C,** Western blot analysis of *Tlr2* RNA and protein extracted from mouse embryonic fibroblasts (MEFs) infected with either non-target pSECC lentivirus (gTom) or Tlr2 targeting pSECC lentivirus (gTlr2). Of note there is no selection cassette in the pSECC plasmid hence residual *Tlr2* expression from non-infected cells.

Supplementary figure 5

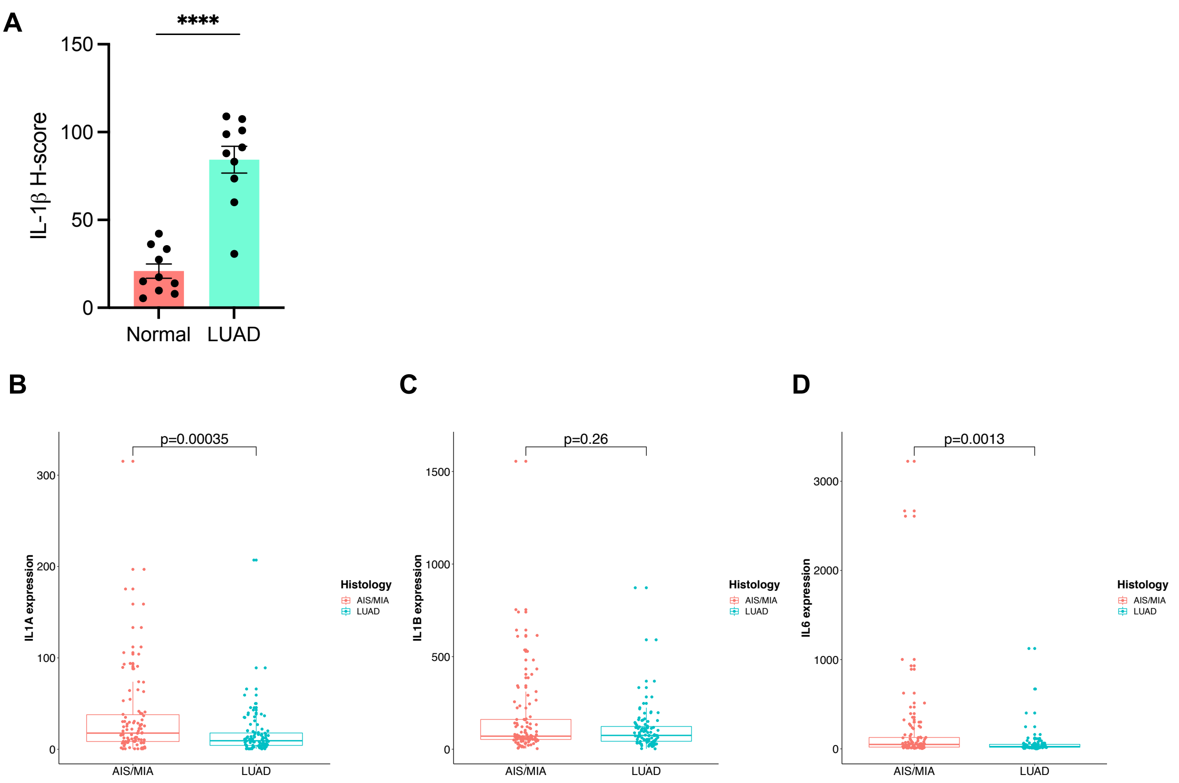


**Supplementary figure 5: A,** Quantification of IL-1*β* IHC in paired normal epithelium (Normal) and lung adenocarcinoma (LUAD). Statistical analysis was performed using the paired students *t*-test. ****p<0.0001. **B,** *IL1A*, **C,** *IL1B* and **D,** *IL6* gene expression was compared between preinvasive (AIS/MIA) and invasive (LUAD) samples. Statistical analysis was performed using the Mann-Whitney test.

Supplementary figure 6

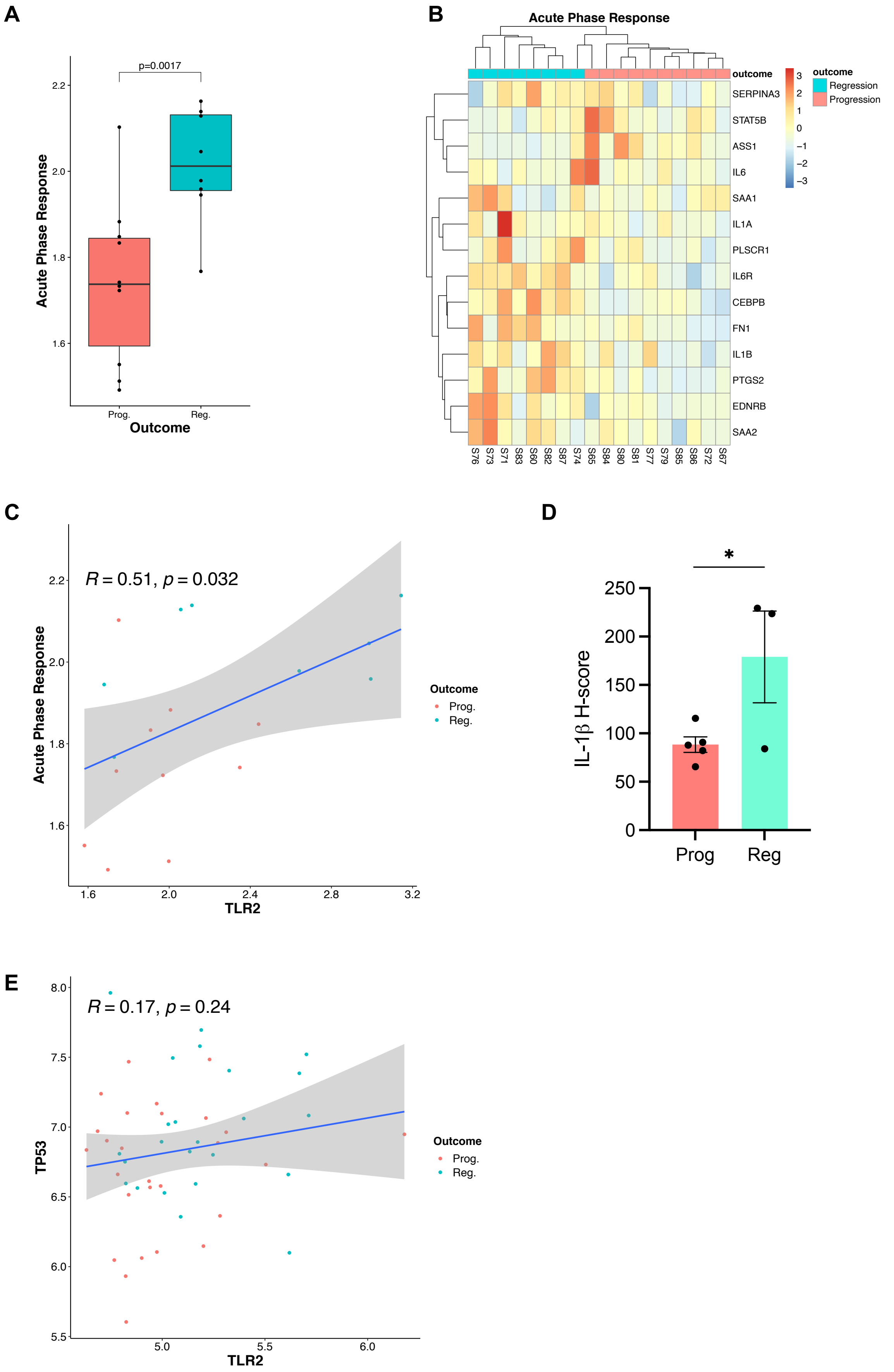

**Supplementary figure 6: A,** Acute phase response gene expression was compared between lesions of equal grade that subsequently progressed to cancer (Prog.) or regressed to normal epithelium (Reg.). Statistical analysis was performed using a linear mixed effects model to account for samples from the same patient. **B,** Heatmap demonstrating Acute phase response gene expression with clear clustering of progressive and regressive lesions. **C,** Scatter plot with Pearson correlation analysis comparing expression of *TLR2* and the Acute phase response expression in progressive (Prog.) and regressive (Reg.) lesions. **D,** IHC quantification IL-1$\beta$ expression in preinvasive LUSC lesions that either progressed to cancer (Prog) or regressed to normal epithelium (Reg). Statistical analysis was performed using the students *t*-test. *$p<0.05$. **E,** Scatter plot with Pearson correlation analysis comparing expression of *TLR2* and *TP53* in progressive (Prog.) and regressive (Reg.) lesions.

#### Supplementary figure 7

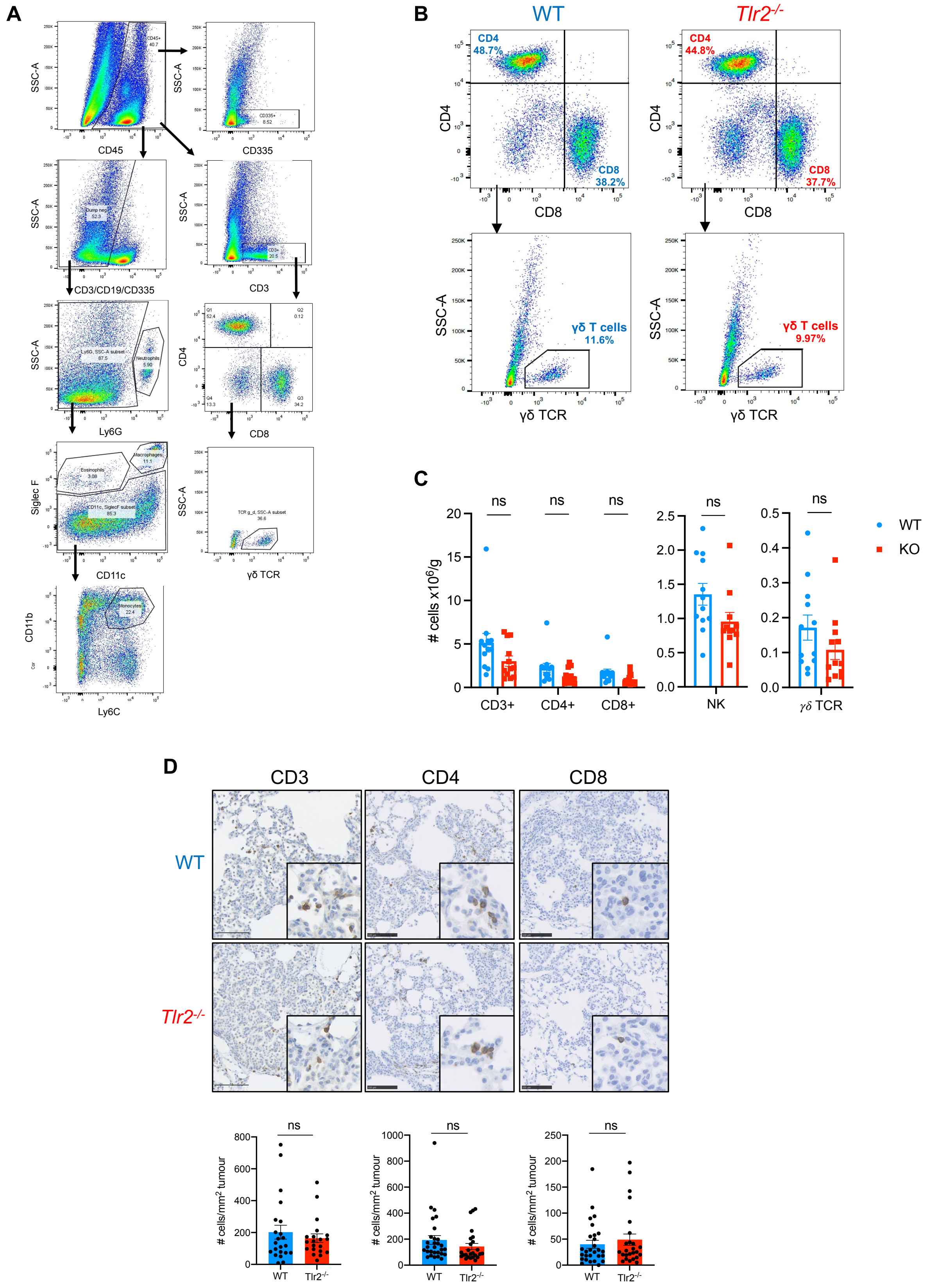

**Supplementary figure 7: A,** Flow cytometry gating strategy used to determine immune cell populations from lung single cell suspensions from tumor bearing mice. **B,** Representative flow cytometry analysis plots of lymphoid populations from whole lung single cell suspensions from WT or *Tlr2*$^{-/-}$ mice. Percentage denotes percentage of parent population. **C,** Corresponding quantification of lymphoid cells from WT (blue) and *Tlr2*$^{-/-}$ (red) mice. CD3+ - total T-cells, CD4+ - CD4 T-cells, CD8+ - CD8 T-cells, NK – natural killer cells, $\gamma\delta$ TCR - $\gamma\delta$ T-cells. n=12 mice per group. **D,** Representative IHC staining for the pan T-cell marker CD3 and specific T-cell markers CD4 and CD8 in WT and *Tlr2*$^{-/-}$ tumors with corresponding quantification. n=5-6 mice per group (five tumors per mouse analyzed). Statistical analysis was performed using the students *t*-test. ns – non-significant.

Supplementary figure 8

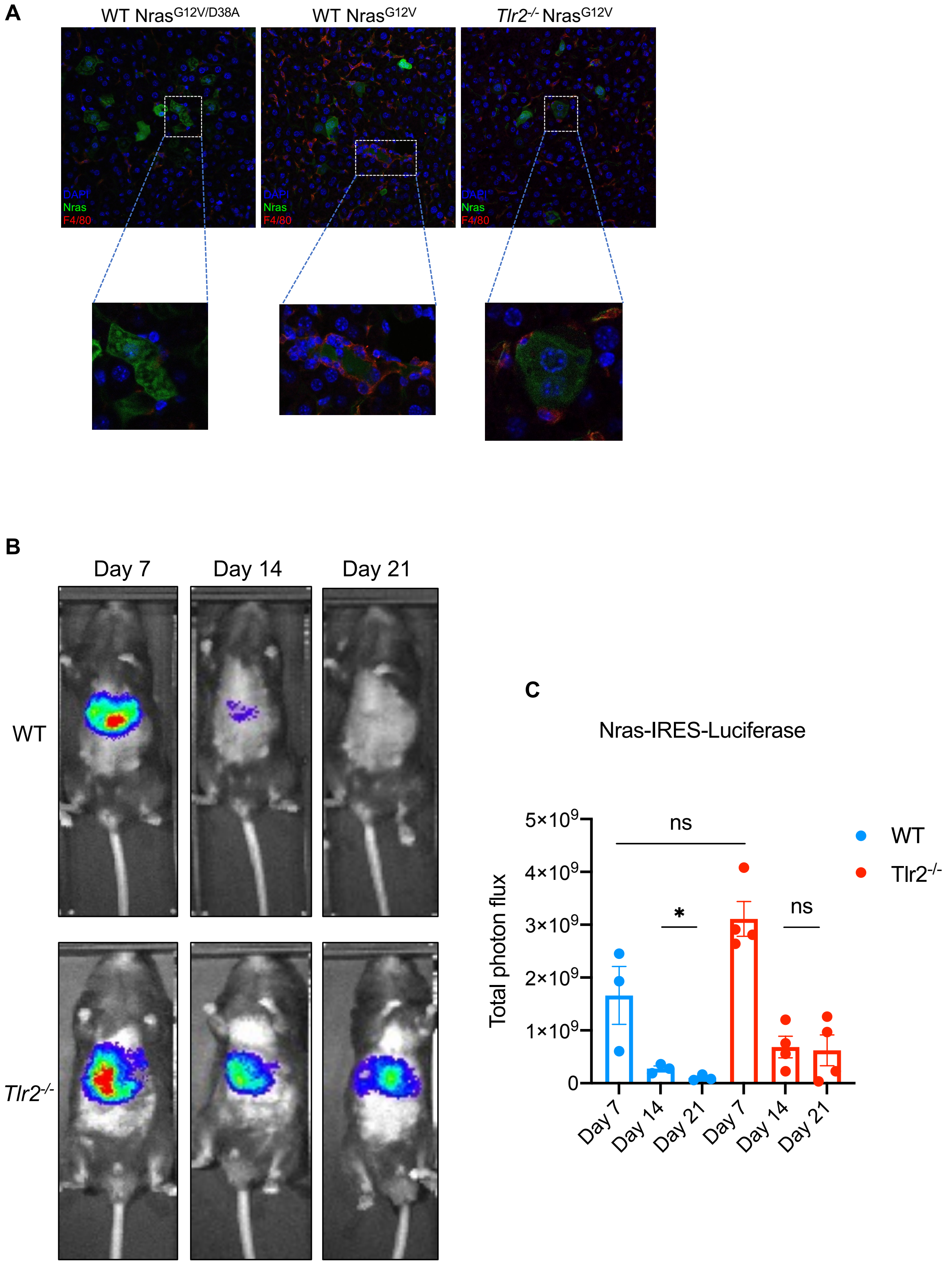

**Supplementary figure 8: A,** Representative co-immunofluorescence staining for DAPI (blue), Nras (green) and F4/80 (red) on liver section from WT or *Tlr2*$^{-/-}$ mice six days after hydrodynamic delivery of oncogenic Nras (*Nras*$^{G12V}$) or negative control (*Nras*$^{G12V/D38A}$) expressing transposons. **B,** Representative bioluminescence images from WT or *Tlr2*$^{-/-}$ mice seven, fourteen and twenty-one days after receiving hydrodynamic delivery of Nras$^{G12V}$-luciferase expressing transposon constructs, with corresponding total photon flux quantification in **C**. Statistical analysis was performed using the students *t*-test. ns – non-significant, *p<0.05.

Supplementary figure 9

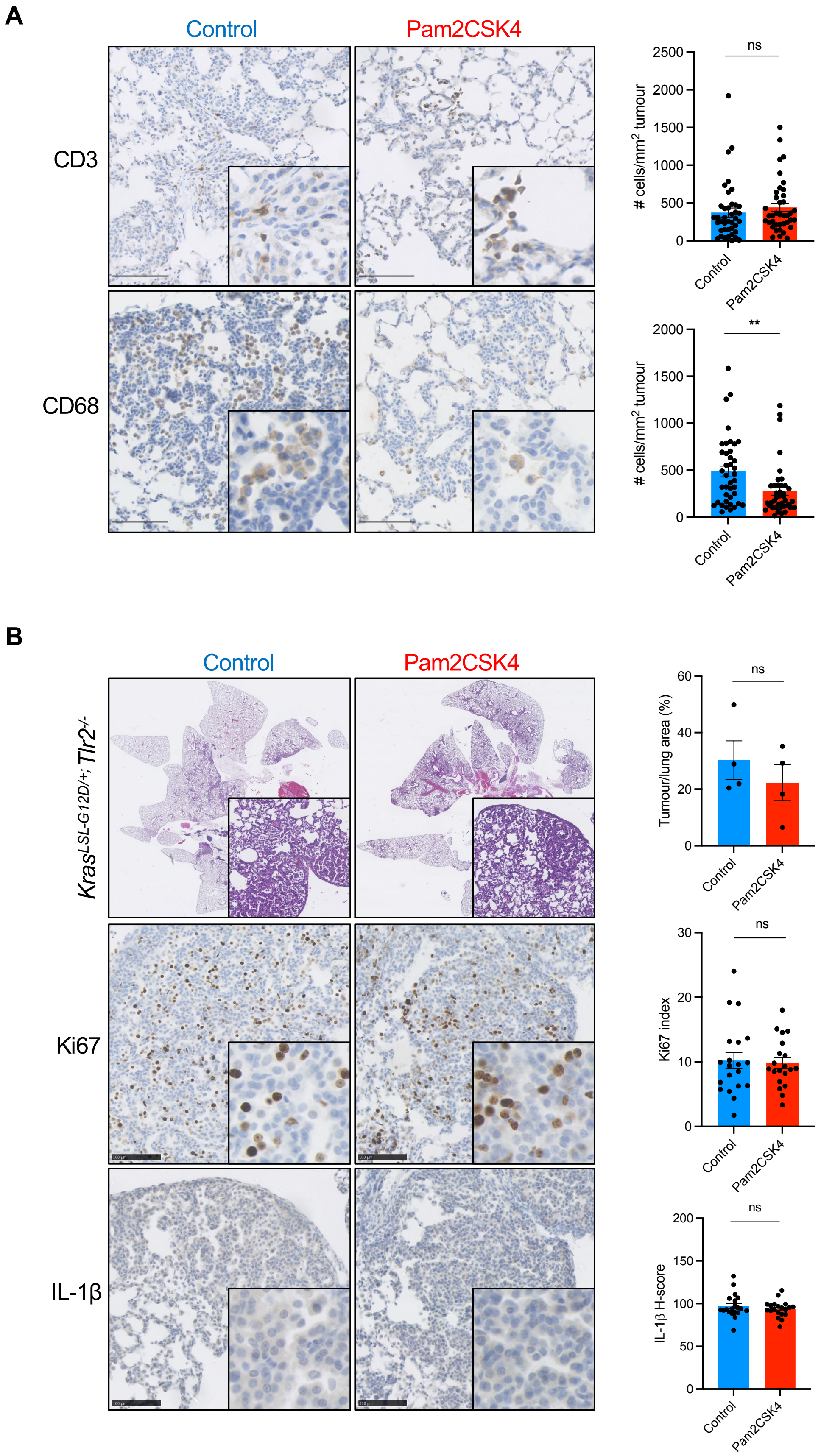


106

**Supplementary figure 9: A,** Representative IHC staining for CD3 and CD68 in lung tumors from *Kras*$^{LSL\text{-}G12D/+}$ mice after either control or Pam2CSK4 treatment, with corresponding quantification. n=8 mice per group (five tumours per mouse analyzed). **B,** Representative H&E and IHC staining for Ki67 and interleukin-1-beta (IL-1$\beta$) in lung tumors from *Tlr2* null Kras$^{LSL\text{-}G12D/+}$ mice (*Kras*$^{LSL\text{-}G12D/+}$*;Tlr2*$^{-/-}$) after either control or Pam2CSK4 treatment, with corresponding quantification. n=4 mice per group (five tumors per mouse analyzed). Statistical analysis was performed using the students *t*-test. ns – non-significant, **p<0.01.

#### Supplementary tables

**Supplementary table 1 – Antibodies used for immunohistochemistry**

| REAGENT or RESOURCE | SOURCE | IDENTIFIER | DILUTION |
|---|---|---|---|
| **Rabbit monoclonal anti-Ki67** | Abcam | Ab16667 | 1:100 |
| **Mouse anti-p21** | BD biosciences | Cat #556431 | 1:200 |
| **Rat anti-p19/Arf** | Abcam | Ab174939 | 1:200 |
| **Goat anti-IL-1 $\alpha$** | R&D systems | AF-400 | 1:50 |
| **Rabbit anti-IL-1 $\beta$** | Santa Cruz | sc-7884 | 1:50 |
| **Rabbit anti-SAA1** | Biorbyt | orb228668 | 1:100 |
| **Mouse anti-Nras** | Santa Cruz | sc-31 | 1:500 |
| **Rat anti-F4/80** | Abcam | Ab6640 | 1:400 |
| **Rabbit anti-CD3** | Abcam | Ab5690 | 1:500 |
| **Rabbit anti-CD4** | Abcam | Ab183685 | 1:1000 |
| **Rabbit anti-CD8** | Abcam | Ab217344 | 1:2000 |
| **Rabbit anti-CD68** | Abcam | Ab125212 | 1:1000 |
| **Rabbit anti-Human TLR2** | Novus Bio | NBP2-24861 | 1:250 |
| **Mouse anti-Human IL1B** | Santa Crus | sc-32294 | 1:100 |
| **Biotinylated goat anti-mouse IgG** | Vector labs | Cat #BA9200 | 1:500 |
| **Biotinylated goat anti-rabbit IgG** | Vector labs | Cat #BA1000 | 1:500 |
| **Biotinylated rabbit anti-goat IgG** | Vector labs | Cat #BA5000 | 1:500 |
| **Anti-rabbit HRP polymer** | Dako | K4003 | N/A |
| **Goat anti-mouse AF488** | ThermoFisher | A11029 | 1:500 |
| **Goat anti-rat AF594** | ThermoFisher | A11007 | 1:500 |

**Supplementary table 2 – Antibodies used for flow cytometry**

| REAGENT or RESOURCE | SOURCE | IDENTIFIER | DILUTION |
|---|---|---|---|
| **Anti-mouse CD45 (Brilliant Violet 421)** | Biolegend | 103133 | 1:100 |
| **Anti-mouse CD3 (PE/Cyanine7)** | Biolegend | 100219 | 1:100 |
| **Anti-mouse CD4 (PE)** | Biolegend | 100407 | 1:100 |
| **Anti-mouse CD8 (Super bright 600)** | Invitrogen | 63-0081-80 | 1:100 |
| **Anti-mouse TCR $\gamma\delta$ (Alexa Fluor® 488)** | Biolegend | 118127 | 1:100 |
| **Anti-mouse/human CD11b (APC)** | Biolegend | 101211 | 1:100 |
| **Anti-mouse CD11c (APC/Cyanine7)** | Biolegend | 117323 | 1:100 |
| **Anti-mouse Siglec-F (PE)** | Biolegend | 155505 | 1:100 |
| **Anti-mouse Ly6C (FITC)** | Biolegend | 128005 | 1:100 |
| **Anti-mouse Ly6G (Alexa Fluor® 700)** | Biolegend | 127621 | 1:100 |
| **Anti-mouse CD19 (PE/Cyanine7)** | Biolegend | 115519 | 1:100 |
| **Anti-mouse CD335 (PE/Cyanine7)** | Biolegend | 137617 | 1:100 |
| **7AAD viability staining solution** | Biolegend | 420403 | 5ul per sample |

**Supplementary table 3 – TLR2 gRNA sequences for pSECC lentiviral constructs**

| **sgRNA ID** | **Forward/Reverse** | **Target sequence** |
|---|---|---|
| gTom | Forward | CACCGGGCCACGAGTTCGAGATCGA |
| | Reverse | AAACTCGATCTCGAACTCGTGGCCC |
| gTlr2 | Forward | CACCGCCTGGAGGTTCGCACACGCT |
| | Reverse | AAACAGCGTGTGCGAACCTCCAGGC |
